# Insulin signaling in pericytes is a master regulator of endocrine and vascular niches in pancreatic islets

**DOI:** 10.64898/2026.11.12.765399

**Authors:** Catarina Andrade Barboza, Luciana Mateus Gonçalves, Roxana Diaz Cruz, Elizabeth Pereira, Ruy Andrade Louzada, Ernesto Bernal Mizrachi, Alejandro Caicedo, Joana Almaça

## Abstract

Insulin plays a crucial role in lowering glycemia, but it is also a powerful anabolic hormone with potent effects on the vasculature. Most studies on insulin vascular actions have focused on endothelial cells. Much less is known about its effects on the mural cells of the microvasculature known as pericytes. This gap in knowledge is particularly relevant for the pancreatic islet microvasculature which is exposed to high insulin levels and relies on local pericytes to efficiently deliver its glucoregulatory hormones. To investigate how insulin affects islet pericyte function and the impact on glucose homeostasis, we generated mice that lack insulin receptors in pericytes (PerIRKO). Loss of insulin signaling in pericytes raised insulin levels, induced islet expansion and beta cell proliferation in the pancreas and in intraocular islet grafts from PerIRKO mice. These changes in beta cell numbers were accompanied by increased islet vascular density, although capillaries were dysfunctional *ex vivo*, compromising islet hormonal responses to an intravenous glucose challenge. Our study supports that insulin itself regulates islet endocrine and vascular niches through previously unrecognized actions on pericytes, revealing a novel mechanism that links the vasculature to endocrine homeostasis.

**Article highlights:**

- Insulin receptors are mostly expressed by vascular cells in mouse and human islets;
- Constitutive knockout of insulin receptors in pericytes (PerIRKO) creates insulin resistance and glucose intolerance;
- Plasma insulin and C-peptide levels are higher in mice lacking insulin receptors in pericytes during feeding but not upon stimulation;
- Increased beta cell proliferation and islet expansion occurs in the pancreas and in intraocular islet grafts from PerIRKO mice;
- Knockout of insulin receptors in pericytes impacts vasomotor responses *ex vivo* interfering with islet hormonal responses to an intravenous glucose challenge.

**Graphical abstract:** 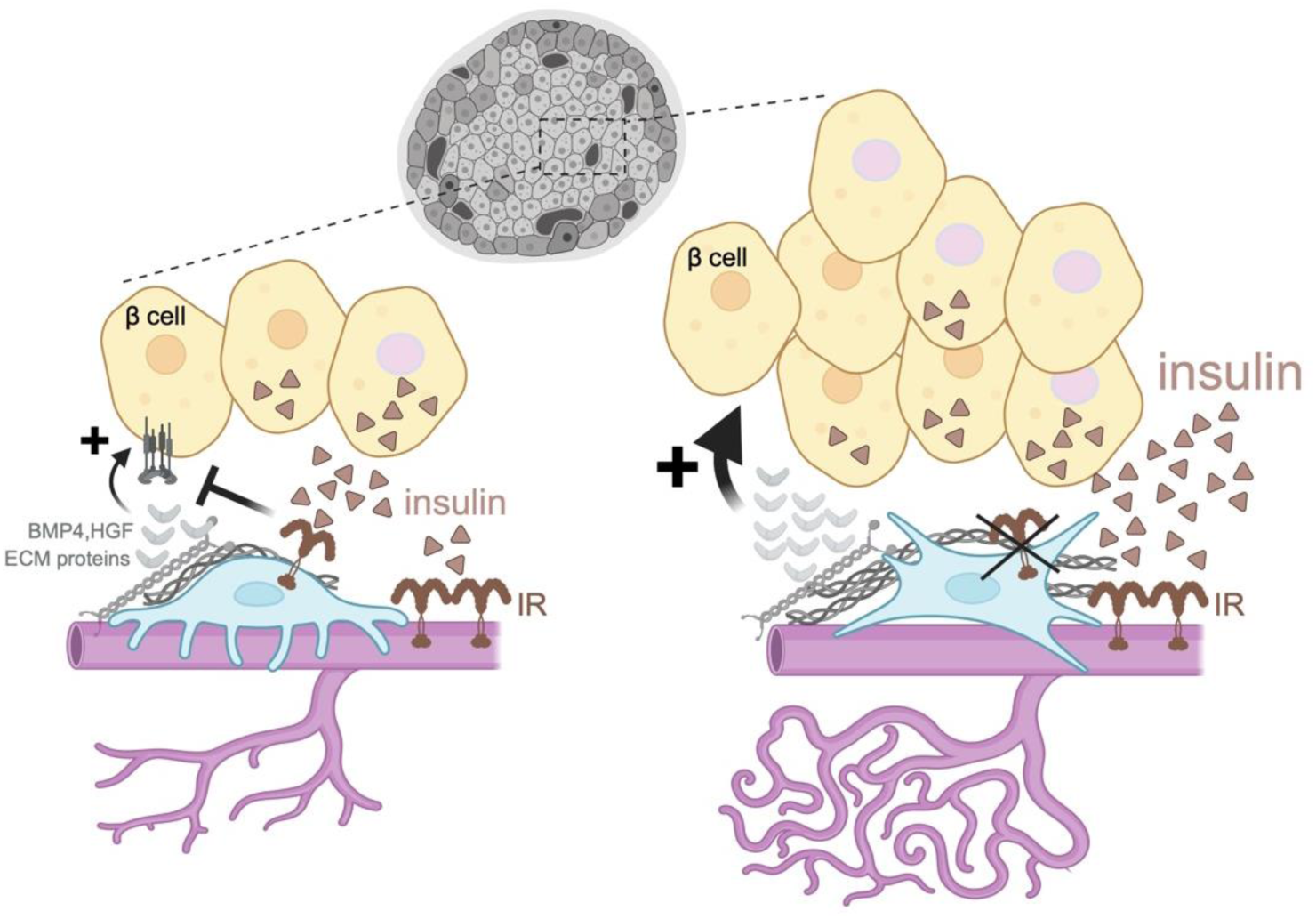

**Graphical abstract and working model:** In pancreatic islets, pericytes have both vascular and trophic functions: not only they stabilize the vasculature (magenta) and control blood flow, but they also produce trophic factors (e.g. BMP4, HGF, ECM components) that stimulate beta cell (yellow) proliferation, maturation and insulin secretion. In this study, we show that islet pericytes express functional insulin receptors (IR) whose deletion, either since birth or in adulthood, leads to beta cell proliferation and islet hypervascularization. In our model, insulin provides negative feedback to islet pericytes to limit the trophic input they provide to beta cells and to stabilize the vasculature. Interfering with insulin signaling in pericytes disturbs this equilibrium and alters pericytes’ trophic and vascular support, altering whole islet homeostasis.

## Introduction

Insulin is known as a glycemia-regulating hormone, but it has broader pleiotropic roles in our body. Abundant research has shown that diseases associated with altered insulin production and action such as diabetes and obesity are characterized by altered vascular responses to vasoactive substances and impaired auto-regulation of blood flow (1). Insulin has potent vascular effects throughout the body (2). Both cellular components of the microvasculature - endothelial cells and pericytes - express functional insulin receptors (3–5). While the role of insulin signaling in endothelial cells has been studied in detail (6–9), much less attention has been given to how insulin can modulate pericyte function in different capillary beds. This is somehow surprising as insulin is even more efficient in enhancing proliferation of pericytes than of endothelial cells (3). Insulin has also been shown to protect pericyte-covered retinal capillaries from ischemia-induced cell death (10) and human brain pericytes from the toxic effects of beta amyloid (11). Acute insulin administration hyperpolarizes retinal pericytes (12), and chronic interference with this signaling pathway leads to retinal hypervascularization (13). Importantly, unlike the inactivation of insulin receptors in endothelial cells which has no major impact on glucose homeostasis under basal conditions (7), constitutive knockout of insulin receptors in pericytes and perivascular fibroblasts creates insulin resistance and hepatosteatosis, preventing weight gain upon high fat diet (14). It has not been explored, however, how insulin signaling in pericytes impacts pancreatic islet function and mass. This is a significant gap in knowledge as beta cells in islets are the source of all the insulin in our body and the islet vasculature is probably exposed to very high levels of this hormone.

Pancreatic islets are highly vascularized mini-organs whose vascular networks consist of capillaries made of endothelial cells covered by pericytes (15, 16). Pericytes act as major regulators of microvascular homeostasis in different tissues (17, 18), but in islets they also actively help to regulate endocrine cell function by displaying both vascular and trophic roles (19). Pericytes impact islet responses to a glucose challenge *in vivo* – by enabling acute changes in islet blood flow that occur upon stimulation with glucose [known as functional hyperemia (20, 21)] and by secreting different trophic factors (such as BMP-4, HGF, NGF) that stimulate beta cell proliferation, maturation and insulin secretion (22–24). Islet pericytes further participate in the production of extracellular matrix (ECM) components (25, 26), which are additional known regulators of beta cell maturation and function (27).

In this manuscript, we hypothesized that pericytes must be able to sense insulin levels to adequately adjust the trophic/vascular support they provide to islet beta cells. To test this hypothesis, we generated two transgenic mouse models that allowed us to interrogate the role of insulin signaling in pericytes early on in development or in adulthood. Our study shows that insulin regulates pericytes’ vascular and trophic functions in islets. Interfering with this signaling impacts glucose homeostasis by affecting islet endocrine and vascular niches: it enhances beta cell proliferation and leads to more vessels in islets, but capillaries are dysfunctional when pericytes lack insulin receptors interfering with islet hormonal responses to an intravenous glucose challenge. Our data show that knocking out insulin receptors in pericytes mimics some of changes that occur in islets in an insulin resistant state such as beta cell and vascular expansions. In summary, our study introduces pericyte insulin signaling as a novel axis in the maintenance of pancreatic islet homeostasis.

## Results

### Insulin receptors are expressed by the microvasculature in mouse and human islets, including endothelial cells and pericytes

Insulin is a pleiotropic hormone whose receptors are widely distributed in almost all vertebrate tissues. The pancreas is no exception to that, and insulin receptors [its β subunits were visualized; (28)] are present in different cell types in endocrine and exocrine tissue compartments in mouse and human pancreas (Figure 1A). Interestingly, while insulin receptors are readily detected at the plasma membrane of acinar/ductal cells in the exocrine tissue, in endocrine compartments most insulin receptor expression is associated with the islet vasculature in mice and humans (Figures 1B,C). To better characterize vascular insulin receptor expression in islets, we used two antibodies against endothelial cell markers: the pan-endothelial marker PECAM-1 (platelet endothelial cell adhesion molecule-1; also known as CD31) and endomucin, a mucin-like glycoprotein found primarily on the surface of endothelial cells lining venous capillaries, venules and veins, but not on arterial endothelium in adult tissues (29). Insulin receptors are expressed by a subset of both arterial and venous endothelial cells in mouse islets, with 40% colocalizing with the pan-endothelial cell marker CD31 but only 20% colocalizing with the venous endothelial cell marker endomucin (Figures 1E,F).

**Figure 1.**
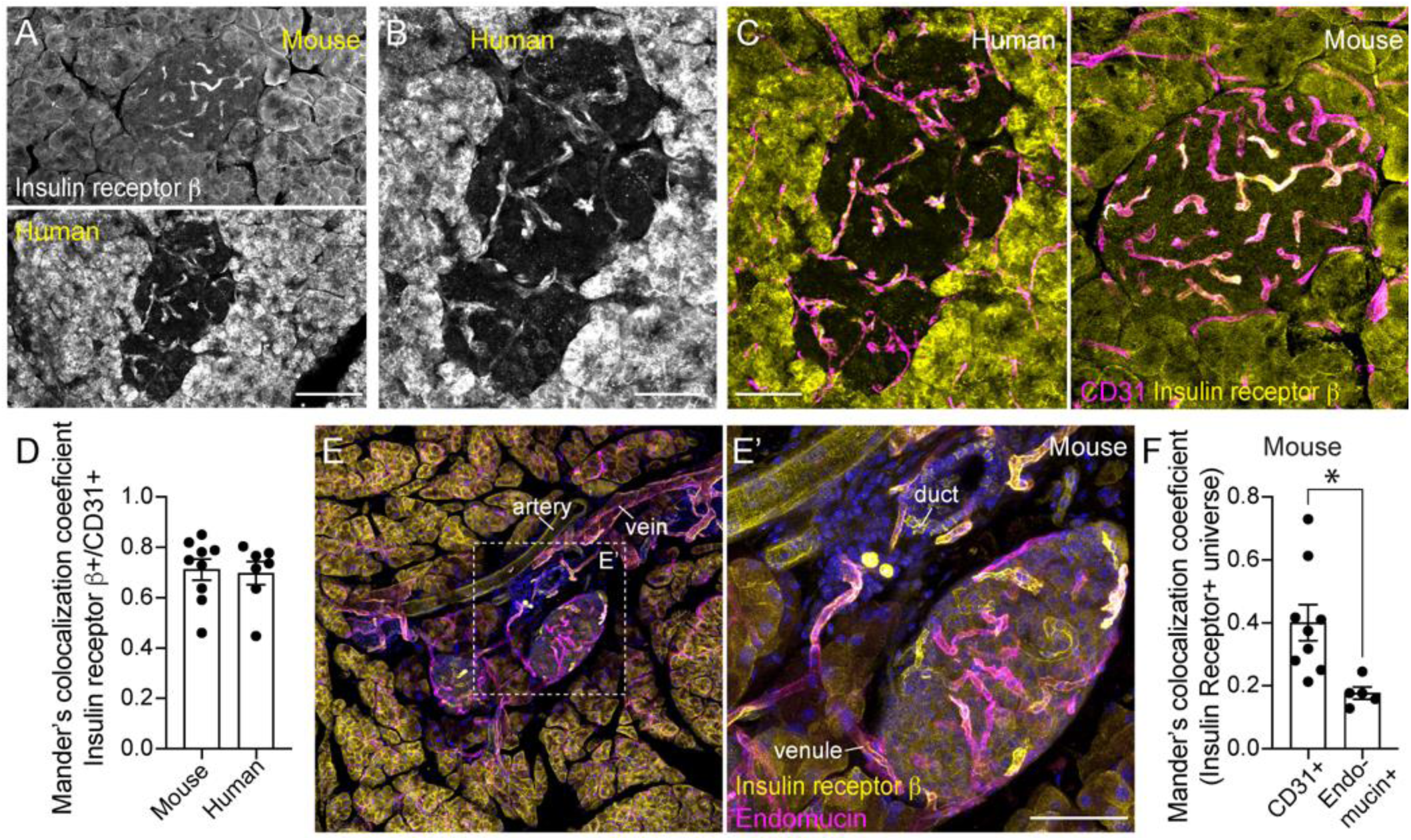
Strong expression of insulin receptors in vascular cells in mouse and human islets. (A) Z-projection of confocal images of mouse (top panel) and human (bottom panel) pancreatic sections immunostained for the insulin receptor β subunit (IRβ chain; gray). (B) Zoom image of the human islet in (A) showing abundant expression of insulin receptors in the vasculature. (C) Insulin receptors (IRβ; yellow) are expressed by endothelial cells (CD31; magenta) in mouse and human islets. (D) Mander’s (M2) coefficient estimating colocalization of insulin receptor β with CD31 in islets from both species (IRβ+ immunostaining /CD31+ universe; n = 7-9 islets/group); (E,E’) Mouse pancreas immunostained for insulin receptors (yellow) and the venous endothelial cell marker endomucin (magenta). (E’) Zoom image of region within dashed square in (E). (F) Mander’s (M1) coefficient as measure of colocalization of insulin receptor with different endothelial cell markers (endomucin or CD31) in mouse islets (n = 9-10 islets/3 mice; \**p* = 0.01 unpaired t-test). Scale bars = 100 μm (A), 50 μm (B,C), 40 μm (E’).

We noticed that, in both mouse and human islets, insulin receptor expression was not equal among all islet vessels (Figures 1C,2A). Colocalization analysis revealed that on average 70% of CD31-labeled endothelium expressed insulin receptors to some extent in islets from both species (Figure 1D). To further characterize which islet capillaries expressed these receptors, we examined their pericyte coverage. Pericytes can be visualized in pancreatic tissues using antibodies against PDGFRβ (platelet-derived growth factor receptor β) or NG2 [neuron-glial antigen 2; (19, 21)]. We noticed that the density of PDGFRβ-labeled pericyte cell bodies was significantly higher around capillaries expressing insulin receptors (Figures 2A,B). In different islets, we could even “guess” which capillaries expressed insulin receptors only by examining their mural cell coverage and creating masks of PDGFRβ receptor staining (Figures 2C and Supplementary Figure 2). High magnification confocal images of vascular niches in mouse and human islets further revealed shared insulin receptor expression by both endothelial cells and pericytes. Although more abundant in endothelial cells (Figure 2D), insulin receptors could be detected around pericytes’ cell bodies and cytoplasmic processes in islets from both species (Figures 2E,F). However, given the very close proximity of pericytes and endothelial cells, which is below the resolution limit of regular confocal microscopes, we further evaluated whether pericytes expressed the insulin receptor gene.

**Figure 2.**
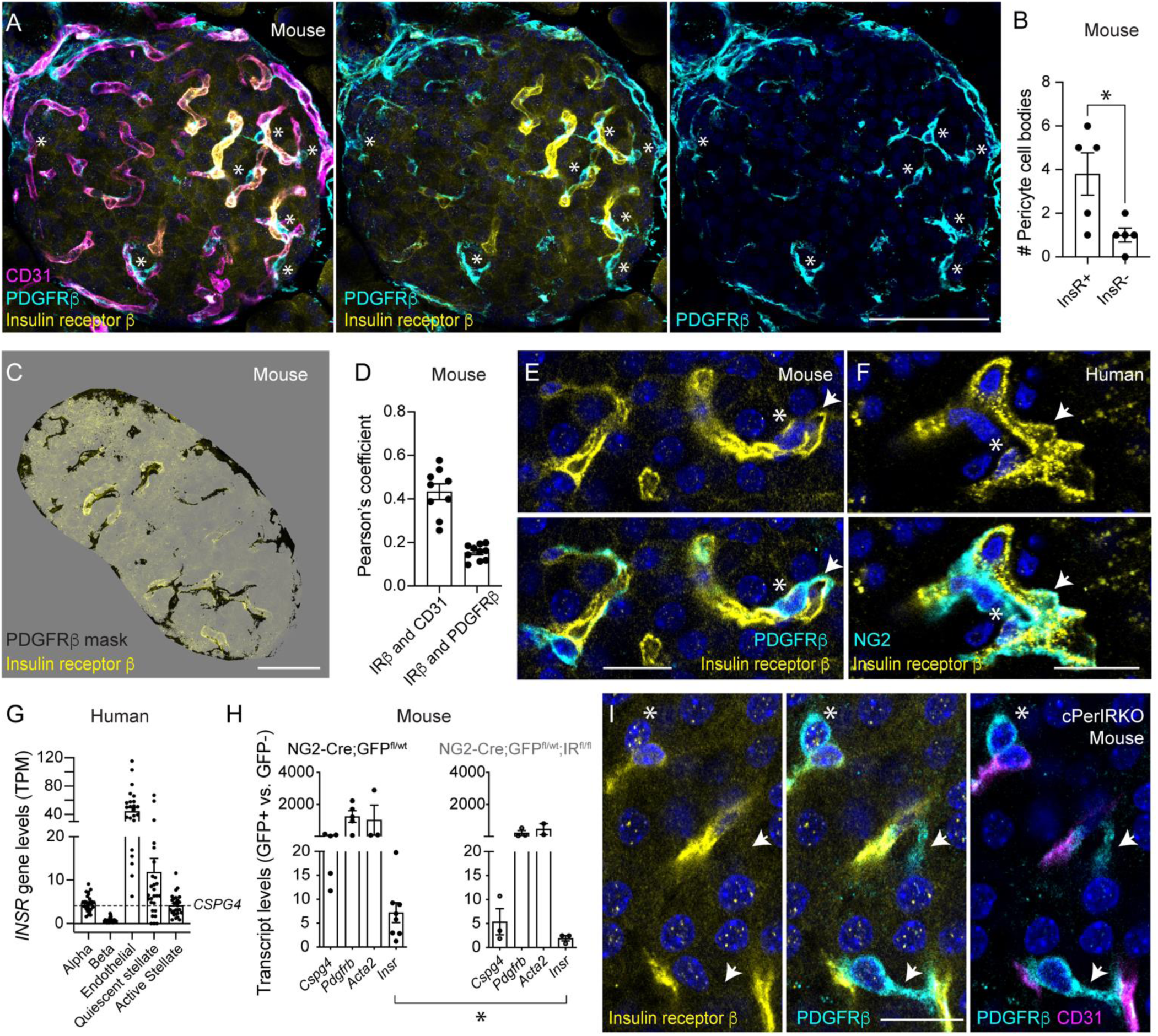
Both endothelial cells and pericytes in the islet express insulin receptors. (A) Z-projections of confocal images of a mouse islet immunostained for the insulin receptor β subunit (IRβ; yellow), endothelial cells (CD31; magenta) and pericytes (PDGFRβ; cyan). Note higher density of PDGFRβ-labeled pericytes (cyan; cell bodies indicated with *) associated with endothelial cells expressing insulin receptors (yellow). (B) Quantification of the number of PDGFRβ-positive pericyte cell bodies covering vessels with and without insulin receptor expression. (C) Mask created using the PDGFRβ signal (black) applied on top of the insulin receptor channel (yellow; image with 40% opacity). Most intra-islet capillaries that express insulin receptor have dense PDGFRβ coverage on their surface. (D) Pearson’s coefficient estimating colocalization between insulin receptor and either CD31 or PDGFRβ in mouse islets (n = 9-10 islets/3 mice). (E,F) Confocal images of vascular niches in mouse (E) and human islets (F) showing pericyte (PDGFRβ or NG2 in cyan; * indicates pericyte cell bodies while arrows indicate cytoplasmic processes) and endothelial cell expression of the insulin receptor (IRβ; yellow). (G) Transcript levels of *INSR* gene in endocrine (alpha and beta) and vascular (endothelial, quiescent and active stellate) cell clusters in human islets from 27 non-diabetic organ donors (data extracted from Dr. Kyle Gaulton’s database: www.isletgenomics.com). Dashed horizontal line indicates the average level of *CSPG4* transcripts in quiescent stellate cells. (H) Pericytes were FACS sorted from islets isolated from mice expressing Cre recombinase under the NG2 (*Cspg4*) promoter/enhancer crossed with floxed-GFP (GCaMP3) reporter mice (control mice) and with IR^fl/fl^ (constitutive PerIRKO). Fold change (2^-ΔΔCt^) in the expression of different genes (pericyte markers - *Cspg4, Pdgfrb, Acta2* - and *Insr*), normalized to *Rn18s* levels, in sorted pericytes (GFP+) versus GFP-cells either from control (left panel) or from constitutive PerIRKO mice (right panel). Significant decrease in the expression of *Insr* in pericytes sorted from PerIRKO islets (islets isolated from 3-8 mice; *p* = 0.04 Mann-Whitney test). (I) Confocal image of blood vessels in islets in the pancreas of a constitutive PerIRKO mouse. Insulin receptor (IRβ; yellow) is expressed by islet endothelial cells (CD31; magenta) but not by pericytes (PDGFRβ; cyan; * and arrows). Scale bars = 20 μm (E,F), 50 μm (C), 40 μm (A), 10 μm (I).

We examined insulin receptor transcript levels in pericyte-enriched clusters derived from mouse and human islets. For that, we isolated islets from mice that expressed a fluorescent reporter (GFP) in pericytes using NG2 (*Cspg4*) Cre mice (20, 26). Real-time qPCR analysis of FACS-sorted islet pericytes revealed detectable levels of the insulin receptor gene (*Insr*) in this cell population, although levels were lower than those of different pericyte marker genes such as *Cspg4*, *Pdgrb* and *Acta2* (Figure 2H). In contrast, gene expression analysis of human islet quiescent sellate cell clusters (which include pericytes; (30); data taken from www.isletgenomics.com) revealed higher levels of *INSR* gene than of the *CSPG4* (which encodes NG2) in this cell population (Figure 2G). Indeed, in comparison with islet endocrine cells (alpha or beta) or with other vascular cells (e.g. active stellate), endothelial and quiescent stellate cells were among those with the highest levels of *INSR* gene in human islets from non-diabetic donors (Figure 2G). In summary, the mouse and the human islet microvasculatures are well equipped to sense insulin that is being released from surrounding beta cells.

### Constitutive loss of insulin receptor signaling in pericytes creates insulin resistance and glucose intolerance

The impact of insulin signaling in endothelial cells on glucose homeostasis had been previously examined and had no major effects on glucose homeostasis under basal conditions (7, 31). To examine the impact of insulin signaling in pericytes, we generated mice that lacked insulin receptors in mural cells by crossing NG2-Cre females with homozygous male mice harboring floxed alleles of *Insr* [(32); constitutive PerIRKO]. Real-time qPCR analysis revelead a ∼75% reduction in *Insr* transcript levels in pericytes sorted from islets isolated from PerIRKO mice (Figure 2H). Insulin receptors were still expressed by islet endothelial cells in constitutive PerIRKO mice but they were not detected around pericyte cell bodies or cytoplasmic processes (Figure 2I).

Constitutive PerIRKO mice were born at the expected Mendelian ratio and by 16 weeks of age there were no significant changes in blood glucose levels in the fed state (Figure 3A; although non-fasting glycemia was lower for control female mice). Upon overnight fasting, hyperglycemia developed in female mice (Figure 3D). Both male and female PerIRKO mice exhibited significantly higher plasma insulin levels but only in the fed state (Figures 3B,C). Intraperitoneal glucose tolerance tests revealed impaired glucose handling in male and female PerIRKO mice (Figures 3D-F), which were also marked insulin resistant as shown in insulin tolerance tests (Figure 3G). These data indicate that insulin signaling in pericytes influences whole-body glucose homeostasis in line with previous findings using a different Cre line which targets pericytes and perivascular fibroblasts during development [*Pdgrfb*-Cre; (14)].

**Figure 3.**
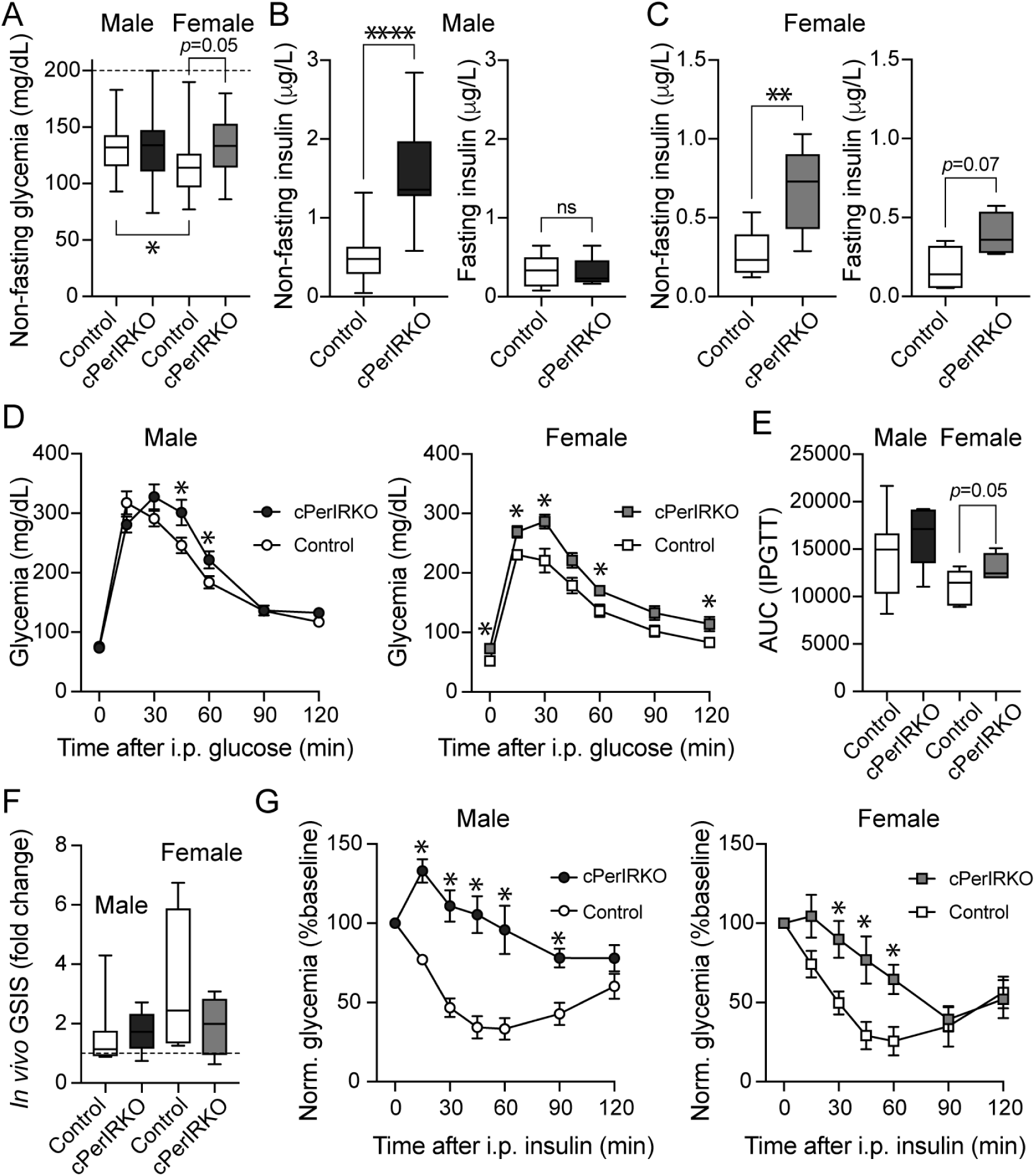
Constitutive loss of insulin receptor signaling in pericytes creates insulin resistance and glucose intolerance. (A) Non-fasting glycemia values in adult (4-month-old) male and female PerIRKO and control mice (n = 7-14 male and 4-5 female mice; One-way ANOVA followed by Tukey’s multiple comparison test; \**p* = 0.03). Control mice are IR^fl/fl^ and NG2-Cre^+/-^ mice pooled together. (B,C) Non-fasting and fasting (overnight) plasma insulin levels for male (B) and female (C) control and PerIRKO mice (n = 7-14 male mice and 5-8 female mice; \*\*\*\**p* < 0.0001 and \*\**p* = 0005; unpaired t-tests). (D) Intraperitoneal glucose tolerance tests (IPGTT; dextrose 2 g/kg body weight) performed after overnight fasting on control and PerIRKO female (right panel; square symbols; n = 4-5 mice) and male mice (left panel; circle symbols; n = 7-14 male mice; \**p* < 0.05 Multiple unpaired t-tests). (E) Quantification of the area under the curve (AUC) of IPGTT curves for control and KO mice (n = 7-14 male and 4-5 female mice; One-way ANOVA followed by Tukey’s multiple comparison test). (F) Fold change in the amount of insulin in the plasma 15 min after i.p. injection of glucose as in (D) (n = 4-8 mice for both genders). Dashed line indicates ratio of 1. (G) Insulin tolerance tests (ITT; insulin 0.75 U/kg body weight) performed after 6h fasting on control and PerIRKO female (right panel; square symbols; n = 4-5 mice) and male mice (left panel; circle symbols; n = 7-13 mice; \**p*<0.05 Multiple unpaired t-tests). Glycemia values normalized to baseline (%).

### Conditional loss of insulin receptor signaling in pericytes in adulthood creates transient hyperinsulinemia

To evaluate whether disturbed pericyte insulin receptor signalling also affected glucose metabolism in adult mice, we crossed NG2-CreER females with homozygous males harboring *loxP* sites flanking exon 4 of the insulin receptor gene (to generate inducible PerIRKO mice). Tamoxifen injection of adult (3-4 months old) NG2-CreER mice enables selective targeting of mural cells (pericytes and smooth muscle cells) in peripheral tissues such as the pancreas [Figure 4A; (21)]. When crossed with floxed tdTomato mice, on day 45 after tamoxifen injection, around 60% of pericytes in endocrine and exocrine pancreas regions expressed the reporter, similar to what has been observed in other tissues using the same Cre line (33). Pericytes expressing tdTomato were FACS-sorted from islets and acinar tissue from inducible PerIRKO and control mice and subjected to qPCR analysis. Forty five days after tamoxifen treatment, insulin receptor transcripts were practically absent in pericytes sorted from islets or acinar tissue from inducible PerIRKO mice. As a control, we examined the levels of insulin-like growth factor 1 (IGF-1) receptor gene as these receptors are also present in pericytes in different tissues [e.g. retina; (34, 35)]. Transcript levels of *Igf1r* gene were similar in islet pericytes from control and KO mice (Figure 4B), indicating selective knockout of *Insr* in pancreas pericytes.

**Figure 4.**
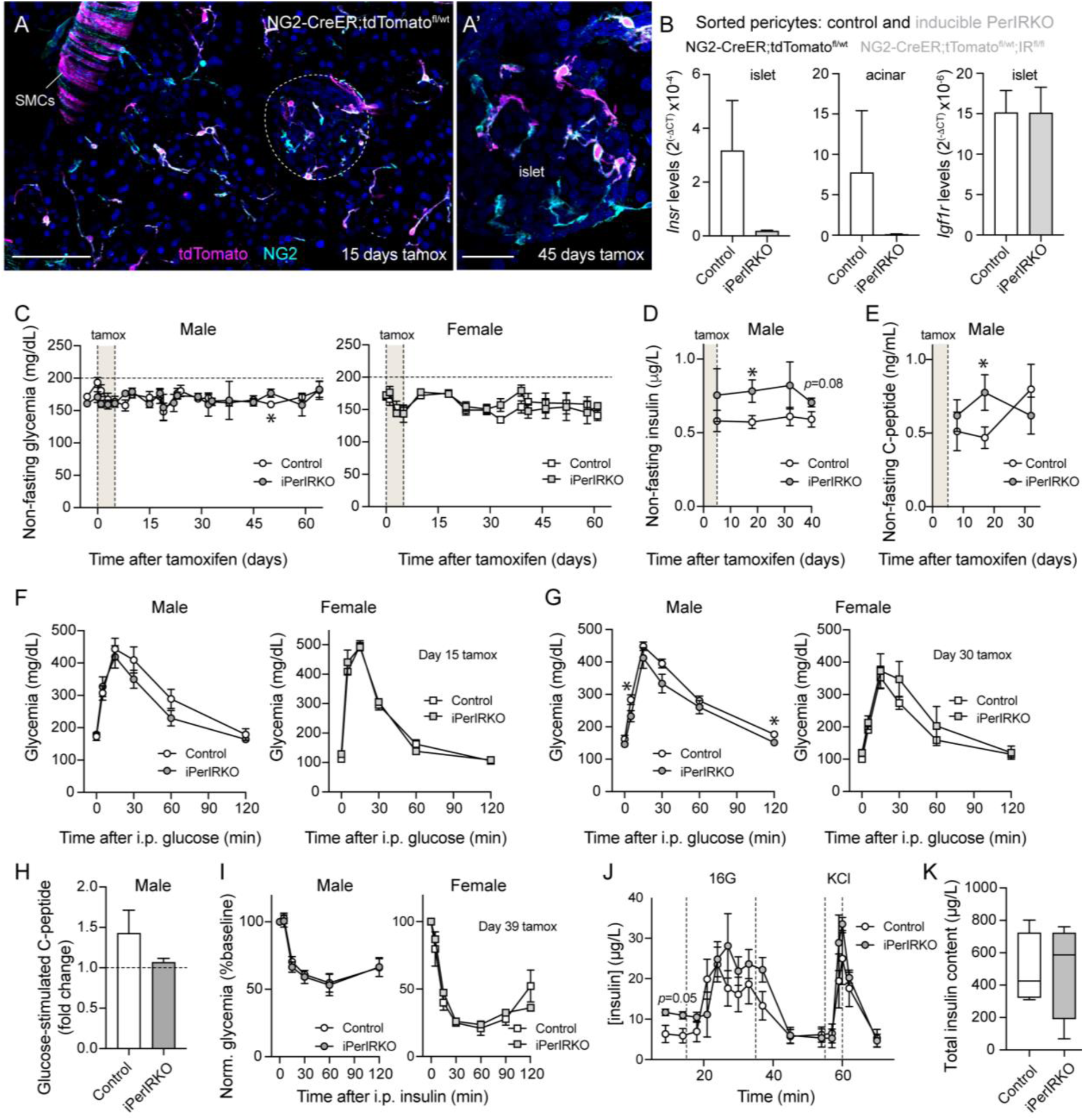
Conditional loss of insulin receptor signaling in pericytes in adult mice leads to a transient hyperinsulinemia in the fed state. (A,A’) Cre recombination is selective and efficient in NG2-CreER mice 15 and 45-days after tamoxifen (on day 45, around 90% of tdTomato-positive cells express NG2 (cyan) and ∼60% of NG2-positive pericytes/SMCs in islets and acinar tissue are tdTomato-positive). Scale bars = 40 μm (A), 20 μm (A’). (B) NG2-CreER^+/-^ female mice were crossed with male IR^fl/fl^;tdTomato^fl/wt^ mice to generate labeled inducible PerIRKO mice. Pericytes were FACS sorted either from islets or acinar tissue from inducible PerIRKO (light gray; n = 3 mice) or control mice (white; n = 4 mice) to examine transcript levels of *Insr* (left and middle panels) and *Igf1r* (right panel). Transcript levels were normalized to those of *Rn18s* in each sample (2^-ΔCt^). (C) Non-fasting glycemia values obtained for adult (3-4-months-old) male and female inducible PerIRKO and control mice up to 60 days after tamoxifen treatment i.p. for 5 consecutive days (light brown rectangle). Control mice include tamoxifen-treated IR^fl/fl^ and vehicle-treated NG2-CreER^+/-^;IR^fl/wt^ mice pooled together. (D,E) Non-fasting plasma insulin (D) and C-peptide levels (E) in inducible male PerIRKO mice at different time points after tamoxifen treatment start (n = 11-13 mice; \**p* < 0.05 unpaired t-tests). (F,G) Intraperitoneal glucose tolerance tests (IPGTT; dextrose 2 g/kg body weight) performed after 6h fasting on control and inducible PerIRKO male and female mice 15 days (F) and 30 days after tamoxifen treatment start (G; n = 7-9 male mice and 3-4 female; \**p* <0.05 multiple unpaired t-tests). (H) Fold change in the amount of C-peptide in the plasma 15 min after i.p. injection of glucose (2g/Kg) conducted 30 days after first tamoxifen injection. Dashed line indicates ratio of 1 (n = 11-13 male mice). (I) Insulin tolerance tests (ITT; insulin 0.75 U/kg body weight) performed after 6h fasting on control and inducible PerIRKO female (right panel; n = 3-4 mice) and male mice (left panel; n = 11-13 mice), 39 days after tamoxifen treatment start. (J) Islets isolated from control and inducible PerIRKO mice were subjected to perifusion experiments (100 islet equivalents per perifusion column). Changes in insulin concentration in perifusate induced by high glucose stimulation (16 mM glucose; 16G) and by KCl depolarization (25 mM). Islets were isolated from 4-5 male and female mice/group (multiple unpaired t-test conducted). (K) Total content of insulin in islets used in each perifusion column (*p* = 0.96 unpaired t-test).

Adult (∼4 months old) male and female inducible PerIRKO (iPerIRKO) and control mice were treated with 5 consecutive doses of tamoxifen and assessed metabolically for up to 60 days after recombination induction. Animals remained normoglycemic and non-fasting glycemia values were similar between the groups (Figure 4C), despite significantly higher fed plasma insulin and C-peptide levels in iPerIRKO male mice already two weeks after tamoxifen treatment (Figures 4D,E). Both male and female iPerIRKO mice remained glucose tolerant, and even a mild improvement was observed for male PerIRKO mice one month after tamoxifen administration (Figures 4F,G). However, while for control mice glucose induced a ∼1.4 fold increase in plasma C-peptide levels 15 min after i.p. injection, C-peptide levels increased only by ∼1.1 fold in male PerIRKO mice (Figure 4H). Animals were equally insulin sensitive as assessed through an insulin tolerance test conducted three weeks after tamoxifen injection (Figure 4I), which contrasted with our findings that constitutive PerIRKO mice were insulin resistant at 4 months of age (Figure 3G).

To explore why non-fasting C-peptide levels were higher in the plasma of inducible PerIRKO (Figure 4E) but did not increase as much upon i.p. glucose stimulation (Figure 4H), we then examined the secretory capacity of beta cells *in vitro* using islets isolated from control and iPerIRKO mice. Stimulating islets with 16 mM glucose or depolarizing endocrine cells with KCl led to similar increases in the amount of insulin released from control and iPerIRKO islets in perifusion experiments (Figure 4J). Total insulin content (100 islet equivalents were used in each perifusion column) did not differ between control and iPerIRKO islets (Figure 4K). In summary, non-fasting plasma insulin (and C-peptide) levels are higher in male mice lacking insulin receptors in pericytes, but not upon stimulation with glucose.

### Knockout of insulin receptors in pericytes leads to islet expansion and beta cell proliferation

After characterizing glucose metabolism, we then extracted the pancreas from mice lacking insulin receptors to examine endocrine cell densities and islet areas. Insulin immunostaining revealed the presence of very large islets in the pancreas of both constitutive and inducible PerIRKO mouse models (Figure 5A). In both models, around 30% of the islets analyzed had insulin-positive areas equal or greater than 40,000 μm^2^, while we did not find such an islet in control mouse pancreata (Figure 5B). The total insulin-positive area in the tissue increased by 4- or 5-fold in the pancreas of inducible and constitutive PerIRKO mice, respectively (Figure 5C). To determine whether the increase in beta cell area resulted from increased proliferation, we stained pancreatic sections from constitutive and inducible PerIRKO mice with an antibody against Ki67. Proliferating (Ki67+) cells could be detected in acinar and ductal compartments in control and KO mouse pancreata, and fewer in endocrine regions (Figure 5D). Indeed, only 5% of the total Ki67+ cell population were beta cells in control mice, but this proportion increased to about 27% and 17% in the pancreas of PerIRKO mice (constitutive and inducible, respectively; Figure 5D). The average percentage of insulin-positive cells that expressed Ki67 went from 0.3% in control islets to ∼1% in both PerIRKO models (Figures 5E,F). In summary, these data indicate that interfering with insulin signaling in pericytes leads to an increase in beta cell proliferation and islet expansion. This increase in islet area was independent of systemic insulin resistance as, unlike constitutive knockout mice, inducible PerIRKO remained insulin sensitive. Interestingly, in both knockout models, besides an increase in area, some islets also lost their circular shapes and typical round morphologies (Supplementary Figure 3).

**Figure 5.**
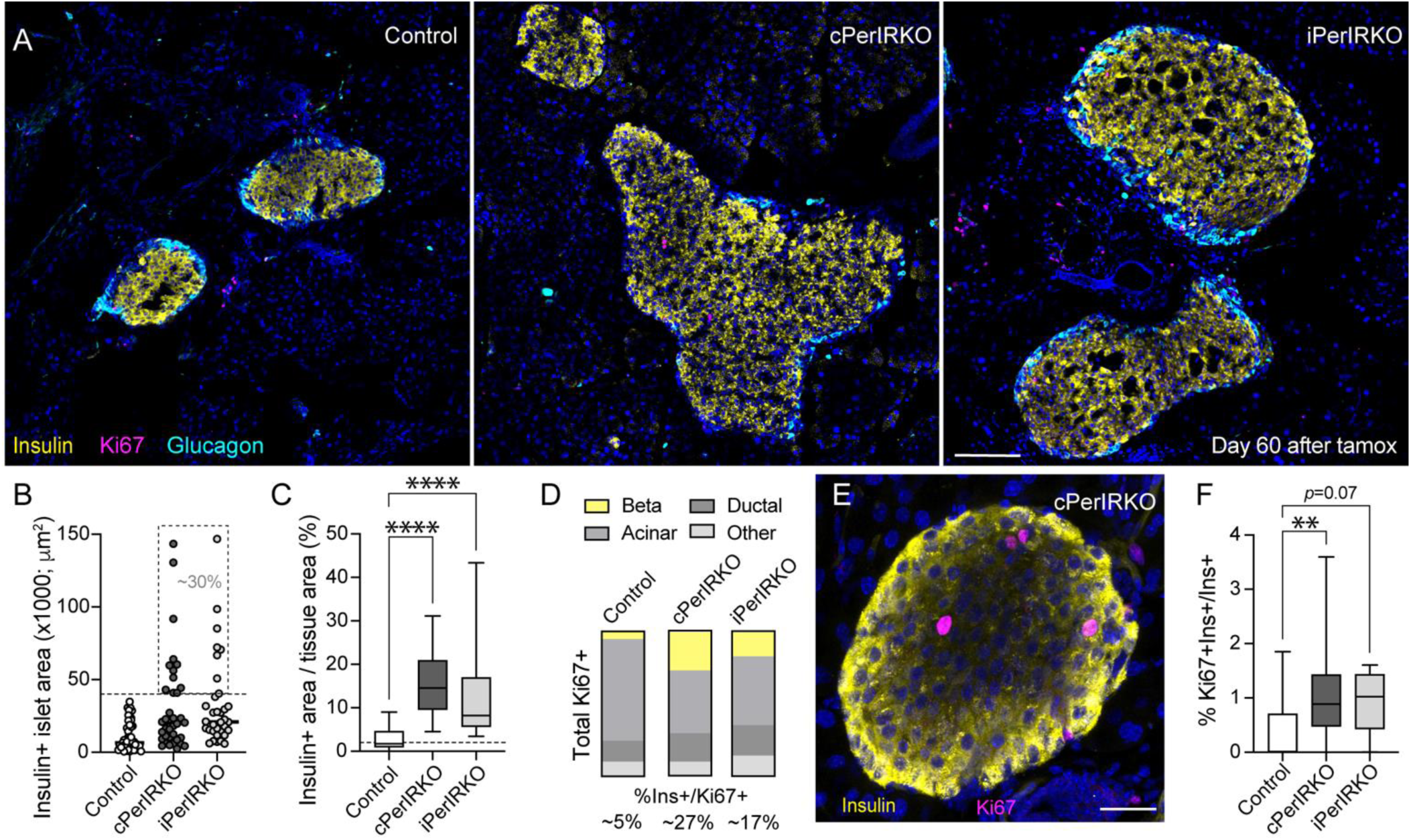
Knockout of insulin receptors in pericytes leads to islet expansion and beta cell proliferation. (A) Z-projection of images of islets in pancreatic sections from control mice (IR^fl/fl^), constitutive (cPerIRKO; 5-months-old) and inducible PerIRKO mice (iPerIRKO; 60 days after tamoxifen treatment) immunostained for insulin (yellow), glucagon (cyan) and the proliferation marker Ki67 (magenta). (B) Insulin-positive areas of islets in the pancreas of control and PerIRKO mouse models (n = 34-47 islets from 3-5 mice). Around 30% of the islets in PerIRKO models had insulin-positive areas equal or greater than 40,000 μm^2^, *versus* 0% in control mouse pancreata. (C) Quantification of the total insulin-positive area divided by the total area of tissue imaged (as %). Dashed line indicates 2% value as beta cells make up 1-2% of the pancreas (n = 13-48 images from pancreases from 3-5 mice; \*\*\*\**p*<0.0001, one-way ANOVA followed by Dunnett’s multiple comparison test). (D) Quantification of the percentage of Ki67-positive cells found in different pancreas compartments of control and PerIRKO mouse models (insulin-positive compartment is shown in yellow and the % of Ins+/Ki67+ is displayed at the bottom). (E) Confocal image of an islet in a pancreatic section from a constitutive PerIRKO mouse immunostained for insulin (yellow) and Ki67 (magenta). (F) Quantification of the percentage of insulin-positive cells that are also Ki67-positive in islets from control and PerIRKO mouse models (n = 8-22 islets from 3-5 mice; \*\**p*=0.003, one-way ANOVA followed by Dunnett’s multiple comparison test). Scale bars = 100 μm (A), 20 μm (E).

### Islet hypervascularity occurs upon knockout of insulin receptors in pericytes

It was surprising to us that the massive increase in islet size and beta cell number found in PerIRKO mice did not translate into better metabolic performance during i.p. glucose tolerance tests. Because the islet microvasculature is also an important regulator of plasma insulin levels and beta cell responses to glucose *in vivo* (36), we then examined vascular trees and their cellular components (endothelial cells and pericytes) in pancreatic sections from inducible and constitutive PerIRKO mice (Figure 6A). Knocking out insulin receptors in pericytes led to a significant increase in the density of CD31-positive endothelial cells in islets, but did not affect the density of NG2 expressing pericytes (Figures 6B,C). Islets from inducible PerIRKO mice not only had more blood vessels but capillaries also looked more tortuous. A subset of them was covered with the matricellular protein periostin (Figure 6D). Periostin is a key player in the development of fibrosis in different tissues such as the lungs (37), which regulates extracellular matrix composition and collagen crosslinking, and it is usually detected only around capillaries at the islet border in control mice (Figure 6F). However, several intra-islet capillaries in inducible PerIRKO, but not in the constitutive model, were densely covered by periostin (Figures 6D,G,I), and intra-islet periostin density showed a fourfold increase (Figure 6E).

**Figure 6.**
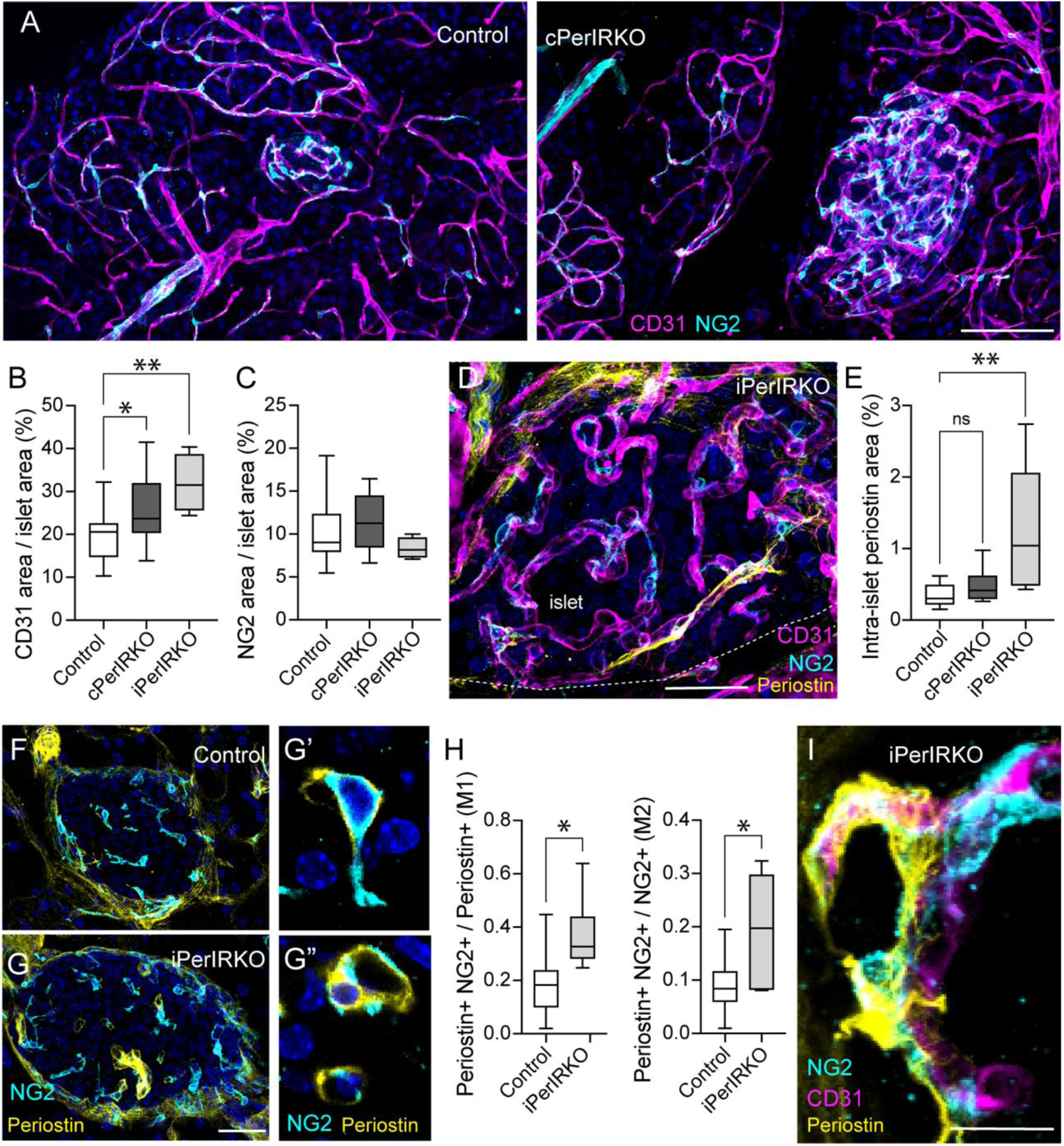
Knockout of insulin receptors in pericytes results in islet hypervascularity. (A) Z-projection of confocal images in fixed pancreatic slices from control (NG2-Cre^+/-^) and constitutive PerIRKO mice immunostained for CD31 (magenta) and NG2 (cyan). (B,C) Quantification of either the total CD31 (B) and NG2 immunostained area (C) divided by the total islet area (as %) for control and both PerIRKO mouse models (n = 7-22 islets from 3-5 mice/group; \**p* = 0.02 and \*\*\**p* = 0.0004, one-way ANOVA followed by Dunnett’s multiple comparison test). For the control group, we pooled data from NG2-Cre^+/-^ vehicle-treated and IR^fl/fl^ mice tamoxifen-treated. (D) Z-projection of confocal images of an islet in an inducible PerIRKO pancreatic section (60 days after tamoxifen) immunostained for CD31 (magenta), NG2 (cyan) and the matricellular protein periostin (yellow). Dashed line denotes islet border. (E) Quantification of intra-islet (excluding the capsule) periostin immunostained area divided by total islet area in sections from control and PerIRKO models (n = 6-8 islets from 3 mice/group; \*\**p* = 0.004, one-way ANOVA followed by Dunnett’s multiple comparison test). (F,G) Z-projection of confocal images of islets from control (tamoxifen-treated) and inducible PerIRKO mice immunostained for periostin (yellow) and NG2 (cyan). Periostin accumulates around some iPerIRKO islet capillaries. (G’,G”) Zoom confocal images of pericytes from the iPerIRKO islet shown in (G) showing NG2 and periostin expression. (H) Mander’s (M1 and M2) coefficients reflecting the extent of colocalization of periostin and NG2 inside islets in control and inducible PerIRKO mice (60 days after tamoxifen; n = 6-8 islets from 3 mice/group; \**p*<0.05, unpaired t-test). (I) Z-projection of confocal images of a pericyte (cyan) covered capillary (magenta) in an islet from an inducible PerIRKO mouse with periostin deposits (yellow) on the surface of the pericyte’s cell body and along its cytoplasmic processes. Scale bars = 40 μm (A,D,F,G) and 10 μm (I).

We then examined whether pericytes could be a potential cellular source of periostin in islets from inducible PerIRKO mice as we had previously reported they are a cellular source of myofibroblasts in islets and contribute to fibrosis (26). We noticed that several islet pericytes expressed periostin (Figure 6G), and the degree of colocalization between periostin and NG2 was significantly higher in islets from inducible PerIRKO mice (Figure 6H). Periostin aggregates could be observed wrapping around PerIRKO islet capillaries, accumulating at the surface of some pericyte cell bodies and along their cytoplasmic processes (Figure 6I). In summary, loss of insulin receptor signaling in pericytes is associated with an increase in vascular density in islets, but some of these capillaries may be dysfunctional as they are covered by pericytes with myofibroblast-like phenotypes.

### Islet vasomotor and hormonal responses to glucose are impaired in PerIRKO mice

We then investigated whether blood vessels in islets from PerIRKO mice remained functional. For that, we produced living pancreas slices from control and knockout mice injected intravenously with a fluorescent lectin before sacrifice. Lectin labeling allowed us to visualize vascular trees in endocrine and exocrine pancreas compartments (Figure 7A). We again noticed an increase in the density of islet functional blood vessels in mice lacking insulin receptors in pericytes (Figure 7B). A closer inspection of capillaries revealed that in some islets in inducible PerIRKO mice vessels looked more tortuous and with a “spring” shape than in control (Figure 7C) but islet capillary diameter remained around 4 μm on average. In contrast, in constitutive PerIRKO several islet capillaries appeared dilated (Figure 7D), and the percentage of vessels with diameters equal or greater than 6 μm doubled (Figure 7E).

**Figure 7.**
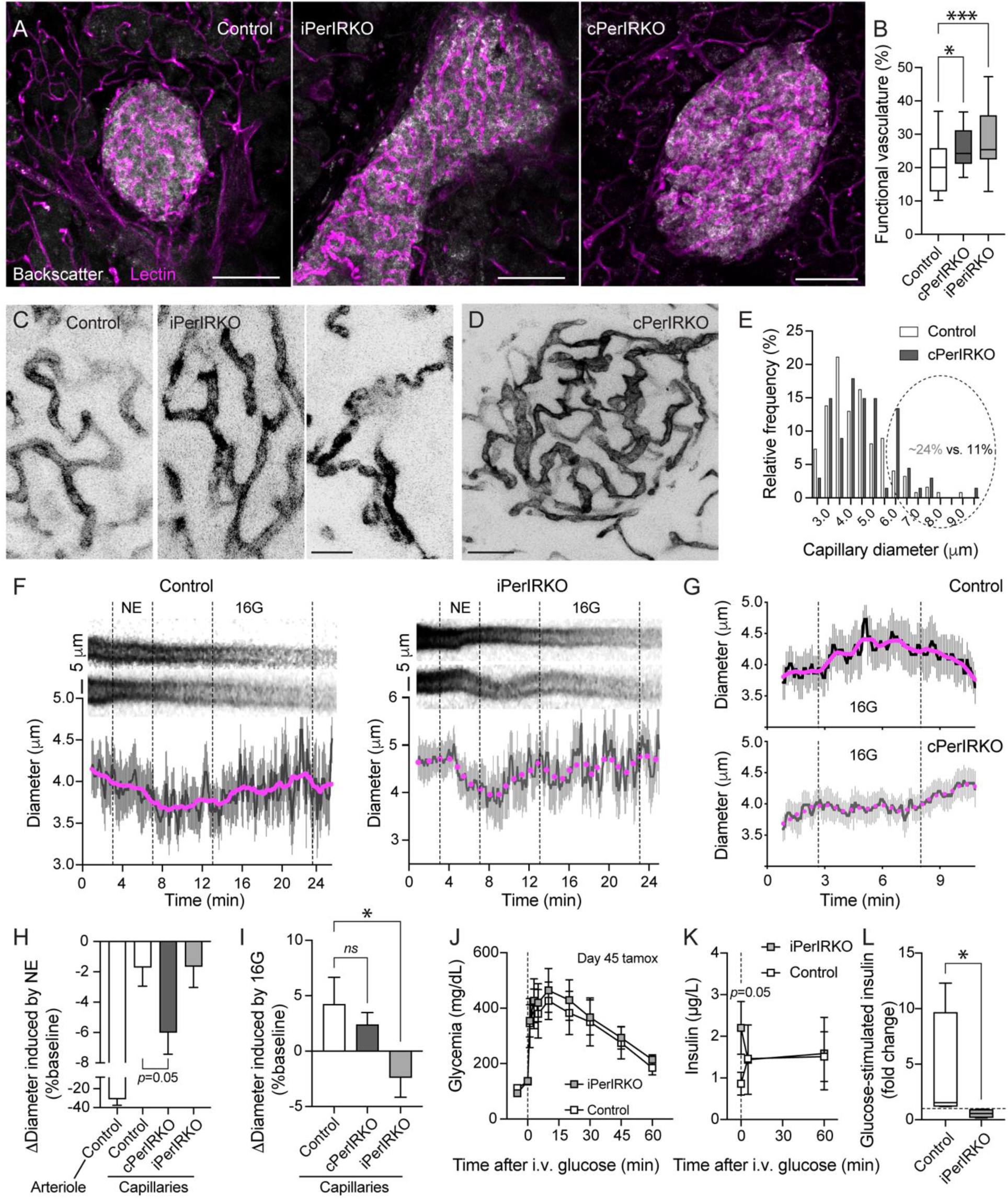
Islet vasomotion and hormonal responses to intravenous glucose are impaired in PerIRKO mice. (A) Living pancreatic slices (150 μm thickness) were produced from PerIRKO mouse models and control mice injected i.v. with a fluorescent lectin (DyLight 649) 10 min before sacrifice. Lectin (magenta) labels vascular networks in the exocrine tissue as well as in islets which can be visualized using the backscatter signal (gray). (B) Quantification of data as in (A), showing the density of lectin-labeled islet blood vessels (functional vasculature) as a percentage of islet area in control and PerIRKO mouse models. Control mice include NG2-Cre^+/-^ and IR^fl/fl^ mice with and without tamoxifen treatment (n=16-33 islets from 3-5 mice/group; * *p* =0.04 and \*\*\**p* = 0.0004, one-way ANOVA followed by Dunnett’s multiple comparison test). (C,D) Inverted grayscale images of lectin-labeled islet capillaries in zoom images (C) or whole islet (D) of control and PerIRKO mouse models. Some capillaries in islets from inducible PerIRKO mice have a strange “spring” shape (right panel in (C)), and some vessels are dilated in islets from constitutive PerIRKO mice (D). (E) Histogram displaying the relative frequency of islet capillary diameters in slices from control and constitutive PerIRKO mice. The percentage of islet capillaries with diameters equal or greater than 6 μm is twofold higher in constitutive PerIRKO than in control. (F) Temporal projections (reslice images; upper panels) and traces (lower panels) showing changes in capillary diameter induced by norepinephrine (NE; 20 μM; applied in 3G) and high glucose (16 mM; 16G). Traces show average ± SEM for 6-8 vessels from control and inducible PerIRKO islets. Smoothened curves of averages of 12 neighbors are shown in pink. (G) Representative traces showing changes in capillary diameter induced by 16 mM glucose for control (upper panel) and constitutive PerIRKO islets (lower panel). Note delay in vessel dilation in PerIRKO mice. Traces show average ± SEM from 6-10 islet capillaries from one islet/genotype. (H,I) Changes in islet vessel diameter (% of baseline) induced by norepinephrine (NE; H) or by 16 mM glucose (I; n = 58-106 capillaries and 15 arterioles, from 4-10 islets from 4-5 mice/group; \**p* = 0.02, one-way ANOVA followed by Dunnett’s multiple comparison test). (J) Intravenous glucose tolerance test conducted after overnight fasting (dextrose injected at 0.75 g/Kg BW) on iPerIRKO and control male and female mice 45 days after tamoxifen treatment. (K) Plasma was collected before, 5 min and 60 min after i.v. glucose injection. (L) Fold change in insulin levels 5 min after glucose injection (n = 4 female mice/group; \**p* = 0.03 Mann-Whitney Test). Dashed horizontal line indicates value of 1. Scale bars = 100 μm (A), 20 μm (C) and 40 μm (D).

We then recorded islet vasomotor responses to stimuli we had previously shown to constrict or dilate islet capillaries such as norepinephrine (NE; 20 μM) and high glucose (16 mM; 16G), respectively (21, 26, 38). Similar to our previous findings, norepinephrine constricted a subset of capillaries in islets from control and both PerIRKO models (Movies S1 and S3; Figures 7F,H; average decrease in capillary diameter of ∼2% *versus* ∼30% reduction for islet arterioles). In some islets from PerIRKO mice, norepinephrine-triggered vasomotion was not characterized by capillary constriction but instead by pulling vessels towards each other like springs (Movie S4). Norepinephrine-induced constriction was even greater for constitutive PerIRKO islet capillaries (average decrease in vessel diameter ∼6%; Figure 7H). In contrast, vasomotor responses to high glucose stimulation were impaired in both knockout mouse models but with some qualitative and quantitative differences: while in control mice high glucose increased average capillary diameter by ∼5%, islet capillaries in inducible PerIRKO decreased their diameters by ∼2% upon high glucose stimulation (Figures 7F,I; Movies S2 and S5). Capillaries in islets from constitutive PerIRKO mice were still capable of dilating upon high glucose but this reponse was delayed and lower in magnitude (average increase in diameter of only 2%) when compared to that of control islets (Figures 7G,I). These data indicate that knocking out insulin receptors in pericytes impacts islet vasomotor responses.

Because islet vasomotor responses to glucose were particularly impaired *ex vivo*, we then examined whether *in vivo* islet responses to intravenous glucose admistration would also be compromised in PerIRKO mice. For that, we subjected inducible PeriRKO mice to an intravenous glucose tolerance test 45 days after tamoxifen injection. Glucose excursion curves were not different between control and iPerIRKO mice (Figure 7J), but insulin levels in the plasma did not increase 5 min after glucose administration in iPerIRKO mice as they did in control mice (Figures 7K,L). These data indicate that knocking out insulin receptors in pericytes interferes with vasomotion and compromises islet hormonal responses to glucose when it is delivered through the endogenous vasculature.

### Beta cell proliferation and vascular remodeling occur in PerIRKO intraocular islet grafts

Given that pericytes are present in many different tissues in the body, we then asked whether manipulating only pericytic insulin signaling in islets was sufficient to induce similar changes in islet endocrine and vascular niches. To answer this question, we isolated islets from adult inducible PerIRKO or control mice before treating them with tamoxifen and transplanted around ∼20 islets into the anterior chamber of the eye of control (Cre-negative;IR^fl/fl^) siblings (Figure 8A). Both male and female mice were used as islet donors and recipients. We imaged longitudinally each eye for 45 days after transplantation and then for 60 days after feeding them a tamoxifen-rich diet. Administering tamoxifen through the diet enables efficient recombination and expression of fluorescent reporters in pericytes in islet grafts from NG2-CreER;tdTomato^fl/fl^ mice (Supplementary Figure 4). For each islet graft, we monitored its size and vascular networks (blood vessels labeled with FITC dextran) as previously done [(39); Figures 8B,C and Supplementary Figure 4]. We noticed that for mice that received control islets, feeding them a tamoxifen diet once islets had fully engrafted did not induce significant changes in graft size or vascular architecture (Supplementary Figure 4; Figures 8B,D). However, for recipients of iPerIRKO islets, tamoxifen diet induced a slight increase in graft area (∼11% increase after 45 days of tamoxifen diet; Figure 8D) as well as a rearrangement of the islet vasculature (Figures 8C,E,F). Vascular rearrangement occurred concomitantly with an increase in the number of cells with backscatter [endocrine cells; (40)] in iPerIRKO islet grafts (Figures 8C’,E’). In eyes with iPerIRKO islets, there were regions at the border of islet grafts that were not visible before tamoxifen diet (yellow rectangle; left panel; Figure 8E) but became well vascularized and with endocrine cells after 45 days of tamoxifen diet (Figures 8E,E’ and Supplementary Figure 4).

**Figure 8.**
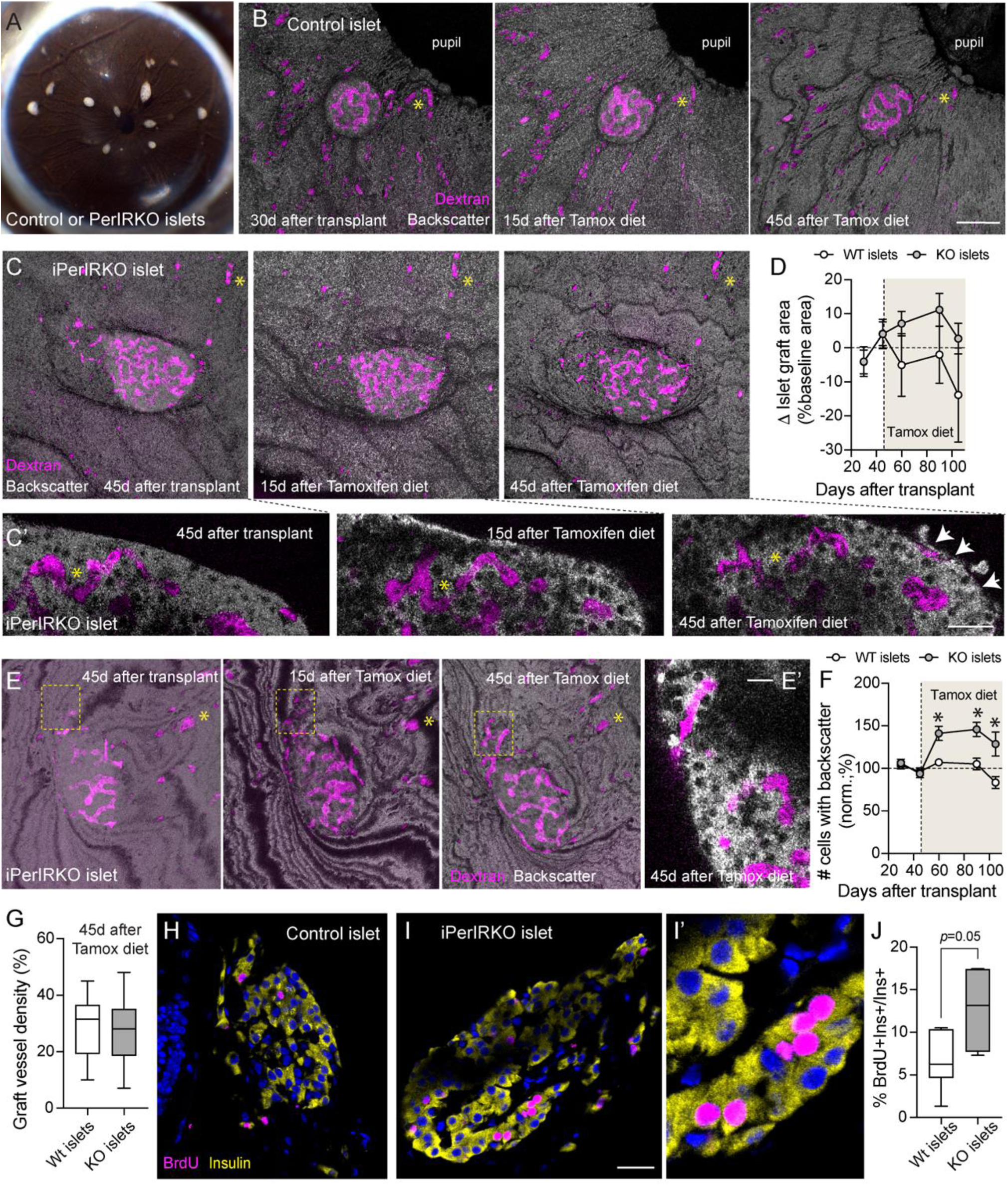
Vascular remodeling and beta cell proliferation in PerIRKO intraocular grafts. (A) Islets were isolated from control (NG2-CreER^+/-^) and inducible PerIRKO mice and transplanted into the anterior chamber of the eye of control (Cre-;IR^fl/fl^) littermates. (B,C) Transplanted eyes with control (B) or inducible PerIRKO (iPerIRKO) islets (C) were imaged longitudinally for 45 days after transplantation and then for additional 60 days while recipient mice were fed a tamoxifen-containing diet. Blood vessels were visualized with FITC-dextran (150 kDa; magenta). Yellow * indicates structures/vessels that were used as reference points for longitudinal imaging. (C’) Confocal images of the same z-plan from the iPerIRKO islet graft shown in (C) at different time points after transplant and tamoxifen diet, showing more cells with backscatter signal in the grafts after feeding a tamoxifen diet (arrows). (D) Quantification of the change in islet graft area over time (baseline = the average area of 2 time points before tamoxifen diet; %) for individual islets from different mice (n = 3 transplanted mice/group). (E,E’) Z-projection of confocal images of an iPerIRKO islet graft at different time points. Dashed yellow rectangle indicates a region in the border of that graft that became vascularized and gained cells with backscatter (E’ zoom confocal image of region within rectangle 45 days after tamoxifen diet). (F) Quantification of change in the number of cells with backscatter in islet grafts from control and iPerIRKO mice at different time points. Shaded area indicates duration of tamoxifen-diet feeding (* p < 0.05; multiple unpaired t-tests). (G) Quantification of dextran-labeled vessel density in control and iPerIRKO grafts 45 days after starting tamoxifen diet feeding. (H,I) At the end of the *in vivo* imaging period, animals received BrdU in the drinking water (1 mg/mL) for two weeks before collecting transplanted eyes for cryosectioning and immunostaining. Confocal images of control (H) and iPerIRKO intraocular grafts (I) in sections of the eye immunostained for insulin (yellow) and BrdU (magenta). Note several beta cells in iPerIRKO islet grafts are BrdU-positive (I’). (J) Quantification of the % of insulin-positive beta cells that are also BrdU-positive in control and iPerIRKO intraocular islets grafts (n = 5 grafts per group; *p* = 0.05 unpaired t-test). Scale bars = 100 μm (B,C), 20 μm (C’,H,I), 10 μm (E’).

At the end of the *in vivo* imaging period (60 days after tamoxifen diet), animals received BrdU, a thymidine analog that labels proliferating cells, in the drinking water for two weeks before collecting transplanted eyes for cryosectioning and immunostaining. Several insulin-positive cells in iPerIRKO islet grafts incorporated BrdU in contrast with few BrdU-positive beta cells in grafts from control mice (Figures 8H-J). In summary, our data indicate that decreasing insulin signaling only in islet pericytes is sufficient to induce beta cell expansion and remodeling of the microvasculature, similar to what we had observed in the native pancreas.

## Discussion

Previous studies examining the impact of insulin on pericytes had focused on peripheral tissues affected in diabetes such as the liver, adipose tissue or the retina (13, 14). However, pancreatic islets are the source of all the body’s insulin but the direct influence of insulin on islet vascular niches had not been examined. Here we investigated how insulin signaling in pericytes impacted pancreatic islet homeostasis and function. Using complementary genetic models that disrupt insulin receptor signaling selectively in pericytes during development or adulthood, we show that loss of pericyte insulin signaling affects both endocrine and vascular islet compartments. These findings establish a previously unrecognized mechanism through which insulin regulates the organ responsible for its own production and output. Rather than acting solely on endocrine cells or on distant metabolic tissues, insulin also acts locally, through pericytes, to coordinate vascular and endocrine niches in islets.

Our initial observation that insulin receptors are mostly expressed by vascular cells within mouse and human islets (Figure 1) prompted us to examine insulin vascular actions in the islet in more detail. We were intrigued that insulin receptor vascular expression was very heterogenous and islet capillaries expressed different levels of these receptors (Figures 1D,2A). In mouse islets, insulin receptor-positive capillaries were not only either arteriolar or venous capillaries (Figure 1E) but had in common their tight association with PDGFRβ-expressing pericytes (Figure 2A). Both insulin and PDGFβ receptors belong to the receptor tyrosine kinase superfamily, and their activation engages similar intracellular signaling pathways. PDGFβ receptors are known to form heterodimers with other tyrosine kinase receptors such as epidermal growth factor receptor (EGFR), and EGFR transactivation is required for PDGF-stimulated cell migration (41). Whether insulin and PDGFβ receptors can also form heterodimers has not been determined, neither whether their transactivation would facilitate signal transduction between endothelial cells/pericytes and vascular actions of both insulin and PDGF at the level of the islet. Pericytes also express IGF-1 receptors in different tissues including the islet (Figure 4B) suggesting they are well equipped to sense growth factors and to adjust blood perfusion to the islet depending on metabolic needs.

Our study shows that, unlike the inactivation of insulin receptors in endothelial cells which had no major impact on glucose homeostasis under basal conditions (7), constitutive deletion of insulin receptors in pericytes created marked insulin resistance in both male and female mice and raised plasma insulin levels (Figure 3). These data are in line with a previous study that had used *Pdgfrb*-Cre mice to knockout insulin receptors in pericytes and perivascular fibroblasts (14). In these animals, systemic insulin resistance developed at 8-weeks of age. The authors reported significant changes in the composition of the liver and adipose tissue, but not of the skeletal muscle. The skeletal muscle is the main site for postprandial glucose disposal (42), and thus a major contributor for whole-body insulin resistance (43). It is thus possible that the lack of insulin receptors in pericytes in the muscle does not affect its composition but instead interferes with capillary recruitment and insulin delivery. Insulin has acute effects on pericytes and it has been shown to hyperpolarize them in the retina, for instance (12). The skeletal muscle vasculature has substantially fewer mural cells than other tissues (44), but pericytes there appear to have very long longitudinal cytoplasmic processes which contact several endothelial cells, forming what has been called a “contineocyte” (45). It is thus possible that, in the skeletal muscle, pericytes mediate acute changes in blood flow induced by insulin, which are needed for proper insulin delivery to muscle tissue. Whether knocking out insulin receptors in pericytes affects insulin-induced changes in muscle blood flow, thus contributing to insulin resistance, will be the focus of future work.

Our study further shows that many of the changes that occur in pancreatic islets in an insulin resistant state – e.g. increased beta cell proliferation and islet mass, increased vessel density and islet capillary dilation, changes in ECM composition, switch in the phenotype of pericytes towards myofibroblasts (46–48) - are mimicked by the knockout of insulin receptors in pericytes. Indeed, interfering with insulin signaling in pericytes creates hyperinsulinemia (Figures 3 and 4), leads to increased beta cell proliferation (Figure 5) and islet functional vessel density (Figures 6 and 7), while simultaneously altering pericytes’ phenotype and impairing microvascular function *ex vivo*. Some of these findings were observed in both the native pancreas and in intraocular islet grafts from knockout mice, revealing a direct role for islet pericyte insulin signaling in coordinating local tissue homeostasis. We have not determined, however, whether impaired insulin signaling in pericytes can interfere with islet adaptive responses during states of higher metabolic demand, contributing to the pathogenesis of diabetes. This will be examined in future studies.

Another important finding from our study is that insulin signaling in pericytes is crucial for proper islet microvascular structure and function (Figure 7). In both PerIRKO mouse models, we noticed differences in islet capillary morphology, diameter and/or function *ex vivo*: in constitutive PerIRKO mice, some islet capillaries are dilated; in inducible PerIRKO, some islet cappilaries are more tortuous, with deposits of the matricellular protein and myofibroblast marker periostin on their surface (Figure 6). We observed a higher degree of colocalization of periostin with the pericyte marker NG2, indicating that pericytes can be turning into myofibroblasts in the absence of insulin signaling and producing this matricellular protein. A switch in the islet pericyte phenotype towards a myofibroblast-like cell could underlie impaired vasodilatory responses to glucose observed in iPerIRKO mice. Indeed, different studies by us and others have reported that altered pericyte contractile phenotype, observed during states of insulin resistance or excessive beta cell mass [e.g. high fat diet feeding (49) or in mice overexpressing a constitutively active Akt in beta cells (26)], is associated with abnormal islet vasomotor responses and blood flow regulation. Because pericyte-mediated acute changes in blood flow are needed for proper islet hormonal responses to glucose in vivo (20), interfering with this acute vascular response could have contributed to lower or defective C-peptide/insulin levels in circulation upon i.p. and i.v. glucose challenges (Figures 4H and 7L). However, the metabolic impact of defective islet vascular responses to glucose could have been masked by the significant expansion in beta cell area we also observe in both PerIRKO mouse models.

In summary, our study supports that pericytes are crucial regulators of endocrine and vascular niches in pancreatic islets, highlighting their role in the maintenance of islet tissue homeostasis. Insulin orchestrates pericyte-mediated control of beta cell proliferation, islet size/shape and vascular density, creating a system that can sense beta cell function and induce changes depending on the output and on metabolic needs. Insulin resistance at the level of the islet vasculature (of pericytes in particular) could thus be a novel cellular/molecular mechanism underlying islet adaptative responses during states of higher demand for insulin, whose failure may contribute to the pathogenesis of diabetes.

## Materials and methods

### Transgenic mouse models

All animal procedures were approved by the Institutional Animal Care and Use Committee (IACUC) at the University of Miami Miller School of Medicine and conducted in strict accordance with NIH guidelines. Animals were housed under controlled conditions (22 ± 2°C, 45-60% humidity, 12-h light/dark cycle) with *ad libitum* access to water and standard rodent chow or specific diets. To generate pericyte-specific insulin receptor knockout mice (PerIRKO) we used transgenic mice that express either Cre recombinase or CreER under the NG2 (*Cspg4*) promoter/enhancer (both strains from Jackson Labs; #008533 and #008538, respectively). The NG2/Cspg4 promoter allows expression or knockdown of genes specifically in mural cells (pericytes and smooth muscle cells) in peripheral tissues such as the pancreas (20, 21, 26). Heterozygous NG2-Cre female mice were crossed with male insulin receptor floxed mice (IR^fl/f^ mice) that harbored LoxP sites within introns 3 and 4 (32) to generate constitutive PerIRKO mice, while NG2-CreER female mice were crossed male IR^fl/fl^ which harbored LoxP sites flanking exon 4 of the insulin receptor gene (from Jackson Labs; #006955) to generate inducible PerIRKO mice. Homozygous IR^fl/fl^ mice were crossed with reporter strains such as tdTomato^fl/fl^ or GFP^fl/fl^ (from Jackson Labs; #007909). For experiments with constitutive PerIRKO, control mice included homozygous IR^fl/fl^ or NG2-Cre^+/-^ littermates, and for those with inducible PerIRKO we used tamoxifen-treated IR^fl/fl^ and vehicle-treated NG2-CreER^+/-^;IR^fl/wt^ mice as control.

### Sex as a biological variable

Because intrinsic sex-differences have been described in murine models of metabolic diseases (50), we used both male and female PerIRKO mice in this study, all as young adults (ages 3-5 months old). Tamoxifen administration did not affect body weight gain in control and inducible PerIRKO male and female mice (Supplementary Figure 1).

### Induction of Cre Recombination

For inducible PerIRKO mice (iPerIRKO) and respective controls (ages 3-5 months old), tamoxifen was administered via two distinct routes. Mice received intraperitoneal (i.p.) tamoxifen injections at a dose of 0.124 mg/g body weight for 5 consecutive days. For mice with intraocular islet grafts from iPerIRKO, Cre recombination was induced via a tamoxifen-supplemented diet (400 mg tamoxifen citrate/kg chow; TD.130860, Inotiv; ∼5g/mouse daily). Dietary treatment was maintained continuously for 2 months from post-transplantation day 45 (as described below).

### Metabolic phenotyping

Systemic glucose homeostasis was evaluated by intraperitoneal glucose tolerance tests (ipGTT) following a 6-h or by an intravenous glucose tolerance test (ivGTT) following an overnight fast. Mice received an i.p. or an i.v. bolus of D-glucose (2 g/kg or 0.75 g/Kg body weight, respectively, in sterile 0.9% NaCl), and tail-vein blood glucose was monitored at 0, 15, 30, 45, 60, and 120 min post-challenge using a handheld glucometer. For insulin tolerance test (ITT), 6h-fasted mice received i.p. regular human insulin (0.75 U/kg body weight), and blood glucose was tracked over 120 min. Non-fasting, fasting (overnight) and upon glucose-stimulation blood samples were collected from the tail vein for quantification of plasma insulin (Ultra-Sensitive Mouse Insulin ELISA kit from Mercodia) and C-peptide levels (mouse C-peptide ELISA kit from Crystal Chem).

### Recording vasomotion in living pancreas slices

Living pancreatic tissue slices were prepared from PerIRKO and control mice injected i.v. with a fluorescent lectin (DyLight 649; 70 μL; #DL-1178-1, Vector Laboratories) 10 min before sacrifice. Briefly, pancreatic tissue pieces (approximately 0.5 cm^3^) were carefully microdissected. The tissue specimens were embedded in 1.5% low-gelling-temperature agarose dissolved in HEPES-buffered solution. Agarose blocks were allowed to solidify at 4°C for 15 min and mounted onto the specimen stage of a vibratome (Leica VT1000S). Slices were sectioned at 150 μm thickness using a slicing speed of 0.1 mm/s and an oscillation amplitude of 1 mm in HEPES-buffered solution containing 0.1% (w/v) BSA. Slices were perfused with HEPES-buffered solution containing 0.1% (w/v) BSA, aprotinin (25 KIU/mL) and 3 mM glucose at 37°C, and imaged on an upright confocal microscope (Leica SP8). Xyzt time-lapse confocal recordings were made (5-second stack intervals over a Z-depth of 50-100 μm). We recorded changes in islet (identified with the backscatter signal) blood vessel diameter induced by glucose (16 mM) and norepinephrine (20 μM; administered in 3 mM glucose). To quantify changes in vessel diameter, we averaged 10 diameter values collected right before and after stimulus admistration, calculated the difference and divided by the baseline (21, 51).

### Islet transplantation into the anterior chamber of the eye and *in vivo* imaging

Pancreatic islets were isolated from male and female adult (3-5 months old) inducible PerIRKO mice and NG2-CreER control mice. Hand-picked islets (∼20 islets per eye) were transplanted into the anterior chamber of the eye of adult, normoglycemic male and female control mice (IR^fl/fl^). Briefly, recipient mice were anesthetized with isoflurane (2.0% in oxygen). Under a stereomicroscope, a small self-sealing corneal incision was made near the limbus. Islets suspended in a minimal volume (∼10-20 μL) of sterile saline solution were aspirated into a custom cannula connected to a Hamilton glass syringe, introduced into the anterior chamber, and deposited onto the vascular bed of the iris.

Islet grafts were imaged biweekly, longitudinally as previously described (39) for 45 days after transplantation to establish a baseline graft size and vascular density. In each imaging session, blood vessels were labeled with a retro-orbital injection of FITC-conjugated dextran (150 kDa; 20 mg/mL in sterile salinesolution, 50 μL per 20 g body weight). To induce pericyte-specific Cre recombination in intraocular islet grafts, mice were placed on a tamoxifen-supplemented diet 45 days post-transplantation. Longitudinal imaging continued biweekly for additional 2 months using a Leica STELLARIS 5 confocal microscope. Mice were anesthetized with isoflurane (2.0% in oxygen) and secured on a custom stereotaxic imaging stage equipped with a thermal pad to maintain body temperature. At the end of this imaging period, animals received BrdU in the drinking water (1 mg/mL) for two weeks before sacrife and eye collection.

### Immunohistochemistry

PerIRKO and control mice were perfused with 4% PFA and their pancreata collected and cryopreserved for immunohistochemistry. Eyes containing islet grafts were enucleated from transplanted mice and fixed in 4% PFA overnight at 4°C. Human pancreatic tissue pieces were fixed overnight in PFA. Tissues were cryoprotected in 30% (w/v) sucrose in PBS, embedded in OCT compound and frozen. Pancreata were cryosectioned (40 μm). Eyes were mounted laterally with the cornea and optic nerve horizontally aligned, and parallel sagittal sections (5 μm) were sequentially cut from the posterior to the anterior chamber. All sections were permeabilized and blocked in 1x Universal Blocking Solution containing 0.3% Triton X-100 for 1h at room temperature. Primary antibodies used include: islet hormones insulin (guinea pig anti-insulin, 1:5, #IR002, Dako) and glucagon (mouse anti-glucagon, 1:1000, #G2654, MilliporeSigma), insulin receptor β subunit (rabbit anti-IRβ, 1:100, #3025, Cell Signaling Technology), endothelial cell markers (rat anti-CD31/PECAM-1, 1:100, #550274, BD Biosciences; rat anti-endomucin, 1:200, #sc-65495, Santa Cruz Biotechnology), pericytes (rabbit anti-NG2/CSPG4, 1:100, #AB5320, MilliporeSigma; goat anti-PDGFRβ, 1:100, #AF385, R&D Systems), extracellular matrix components (rabbit anti-collagen IV, 1:300, #ab6586, Abcam; goat anti-periostin, 1:300, #AF2955, R&D Systems). For cell proliferation assessment, we stained for Ki67 (rabbit anti-Ki67, 1:200, #PA5-19462, Invitrogen) or for BrDU incorporation (rat anti-BrdU, 1:150, #OBT0030G, AbD Serotec) after heat-induced antigen retrieval was performed using a pressure cooker (110°C, 5 min) prior to the blocking step. Species-specific fluorophore-conjugated secondary antibodies (Alexa Fluor 488, 568, 594, 647 and 790; 1:500; Invitrogen) and DAPI (1 μg/mL) were applied for nuclear and target visualization. Confocal Z-stacks were acquired on a Leica STELLARIS laser scanning confocal microscope.

Confocal images of islets in the pancreas were taken every 3-4 μm, spanning around 40 μm of the islet. Z-projections of confocal images were analyzed using ImageJ software to quantify the total area immunostained with CD31, NG2 or periostin divided by the total islet area. For intra-islet periostin density, the region of interest (ROI) drawn around the islet did not include the capsule. Mander’s and Pearson’s colocalization coefficients were calculated to estimate colocalization between insulin receptor β with CD31 or PDGFRβ and between periostin and NG2, using the ImageJ plugin “Just Another Co-localization Plugin” (https://imagej.nih.gov/ij/plugins/track/jacop2.html). Colocalization analysis was performed on 3 different confocal planes for each islet and an average value was calculated for each islet and plotted in the figure.

### Pericyte sorting and gene expression

To isolate pericytes, islets and acinar tissue were isolated from tamoxifen-treated PerIRKO mice crossed with reporter strains (NG2-CreER^+/-^;IR^fl/fl^;tdTomato^+/-^) and control mice (NG2-CreER^+/-^;tdTomato^+/-^) and cultured overnight at 37°C and 5% CO_2_. Islets were dissociated with 1x trypsin at 37°C for 5 min with gentle pipetting. Digestion was stopped with ice-cold HBSS containing 5% FBS, and the resulting cell suspension was filtered through a 35-μm cell strainer. Cells were resuspended in 5% FBS/HBSS containing DAPI (1:10,000) to exclude non-viable cells. Cell sorting was conducted on a FACS Aria II flow cytometer (BD Biosciences) and collected directly into lysis buffer. Total RNA from FACS-sorted pericytes (tdTomato or GFP+) and from other islet cells (tdTomato- or GFP-) was extracted using the RNeasy Micro Kit (Qiagen) according to the manufacturer’s instructions. Reverse transcription was executed using High-Capacity cDNA Reverse Transcription Kit (Applied Biosystems/Thermo Fisher Scientific). Real-time quantitative PCR (RT-qPCR) was performed using TaqMan Gene Expression Assays on a StepOnePlus Real-Time PCR System (Applied Biosystems). We examined changes in the expression of different pericyte genes - *Cspg4, Pdgfrb, Acta2 –* as well as *InsR* and *Igf1r*. *Rn18s* was used as the housekeeping gene.

### Perifusion experiments

Dynamic islet hormone secretion was evaluated by placing hand-picked islets (100 islets per chamber) in a temperature-controlled perifusion system (PERI4; Biorep Technologies, Miami, FL, USA) continuously perfused at a flow rate of 100 μL/min at 37°. Following baseline stabilization, islets were sequentially challenged with high D-glucose (16 mM for 20 min), and depolarizing KCl stimulus (25 mM for 5 min). Perifusate fractions were collected into 96-well plates at 60-second intervals and stored at -20°C until analysis. Hormone levels in supernatant were determined with an Insulin Elisa kit (from Mercodia).

### Statistical analyses

For statistical comparisons we used Prism 10 (GraphPad software) and performed one-way ANOVA followed by Dunnett’s or Tukey’s multiple comparisons tests to compare datasets obtained for control, constitutive and inducible PerIRKO mouse models. We performed unpaired t-tests or Mann-Whitney tests when comparing datasets from control mice with only one PerIRKO mouse model, and multiple unpaired t-tests when comparing datasets over time (e.g. IPGTT and ITT). *p*<0.05 was considered statistically significant. Throughout the manuscript, data are presented as average values for each islet in box-and-whiskers plots or as mean ± SEM when occurring over time.

## Acknowledgements

The authors would like to thank Dr. Elizabeth Soza from the Histology Core and Dr. Oliver Umland from the flow cytometry core at the Diabetes Research Institute, University of Miami Miller School of Medicine. The authors are also very grateful to the assistance of the Pre-clinical Cell Processing and Translational Model Core at the Diabetes Research Institute, under the direction of Dr. Alejandro Tamayo Garcia, for its valuable contributions in metabolic phenotyping of animals in this study. This manuscript was supported by Office of Research Infrastructure Programs of the National Institutes of Health under award number 1S10OD038196-01, for the use of the DRI Stellaris 5 Live-Animal Imaging Confocal system.

## Author contributions

C.A.B and L.M.G performed immunohistochemistry, functional imaging with living pancreas slices, assessment of glucose metabolism in PerIRKO mice, and analyzed data; C.A.B. performed longitudinal imaging of intraocular islet grafts; R.D.C sorted islet pericytes and analyzed gene expression; E.P. genotyped and maintained the animal colony; R.A.L. optimized insulin receptor staining; A.C. transplanted islets into the anterior chamber of the mouse eye; J.A. performed initial experiments with the model, designed the study, analyzed data and wrote the manuscript. All authors had periodic meeting to discuss data, plan experiments, and contributed to the manuscript. Further information and requests for resources or reagents should be directed to the Lead Contact, Joana Almaça.

## Data Access and Responsibility

J.A. is the guarantor of this work and, as such, has full access to all the data included in the study and takes responsibility for the integrity of the data and the accuracy of the analysis. Any requests should be mailed to.

## Data and Resource Availability

All data generated or analyzed during this study are included in the published article (and its online supplementary files). These manuscript used single-cell RNAseq data extracted from Dr. Kyle Gaulton’s publicly available database: www.isletgenomics.com.

