## Supplemental Material for "Insulin signaling in pericytes is a master regulator of endocrine and vascular niches in pancreatic islets"

**Supplementary Material – Figures and Movies**

**Supplementary Figure 1**

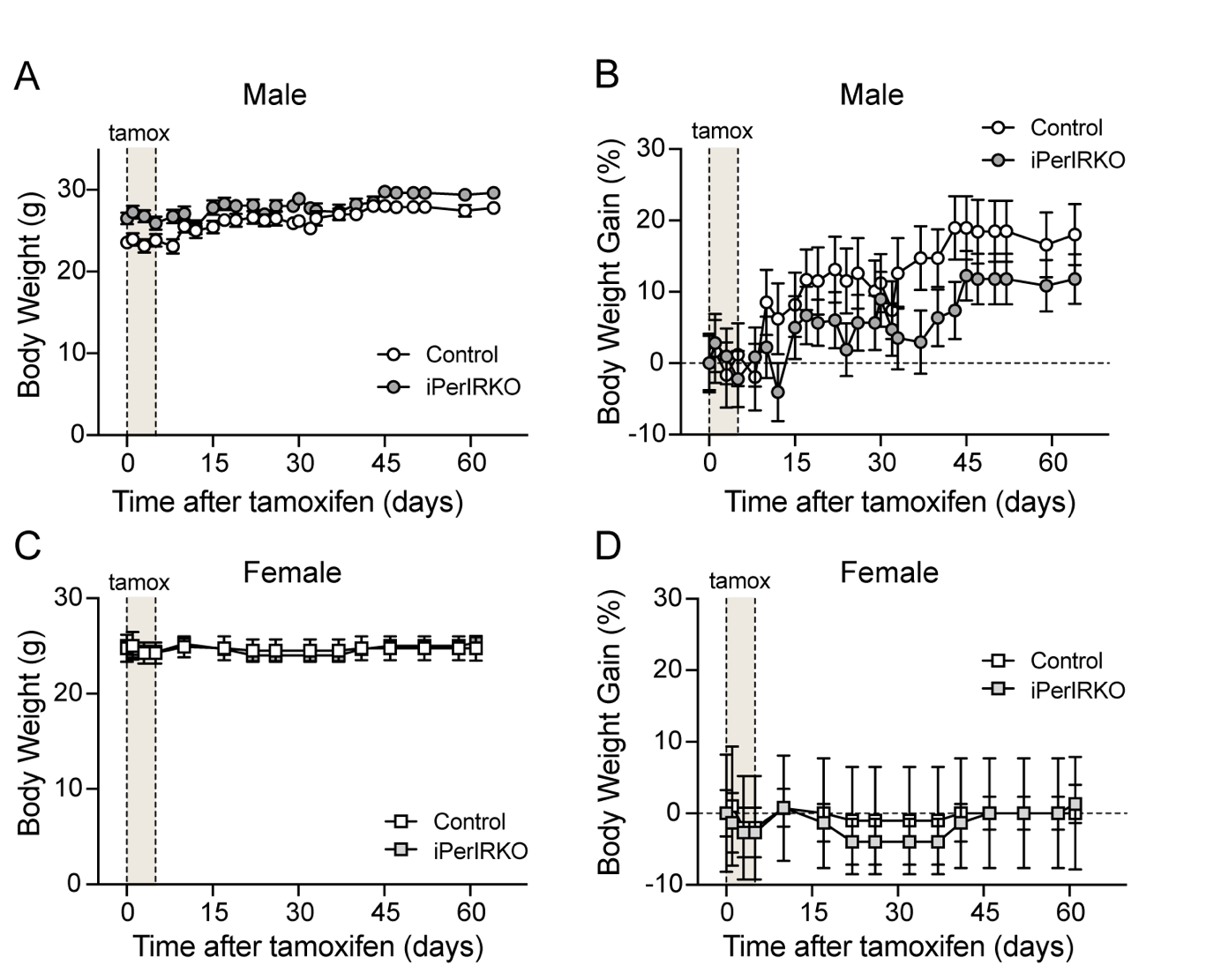

**Supplementary Figure 1. Tamoxifen treatment does not interfere with body weight gain of inducible PerIRKO and control mice.** Male (A,B) and female (C,D) inducible PerIRKO mice (iPerIRKO) and respective controls (ages 3-4 months old) received 5 consecutive intraperitoneal (i.p.) tamoxifen injections at a dose of 0.124 mg/g body weight. Body weight was monitored throughout the study.

**Supplementary Figure 2**

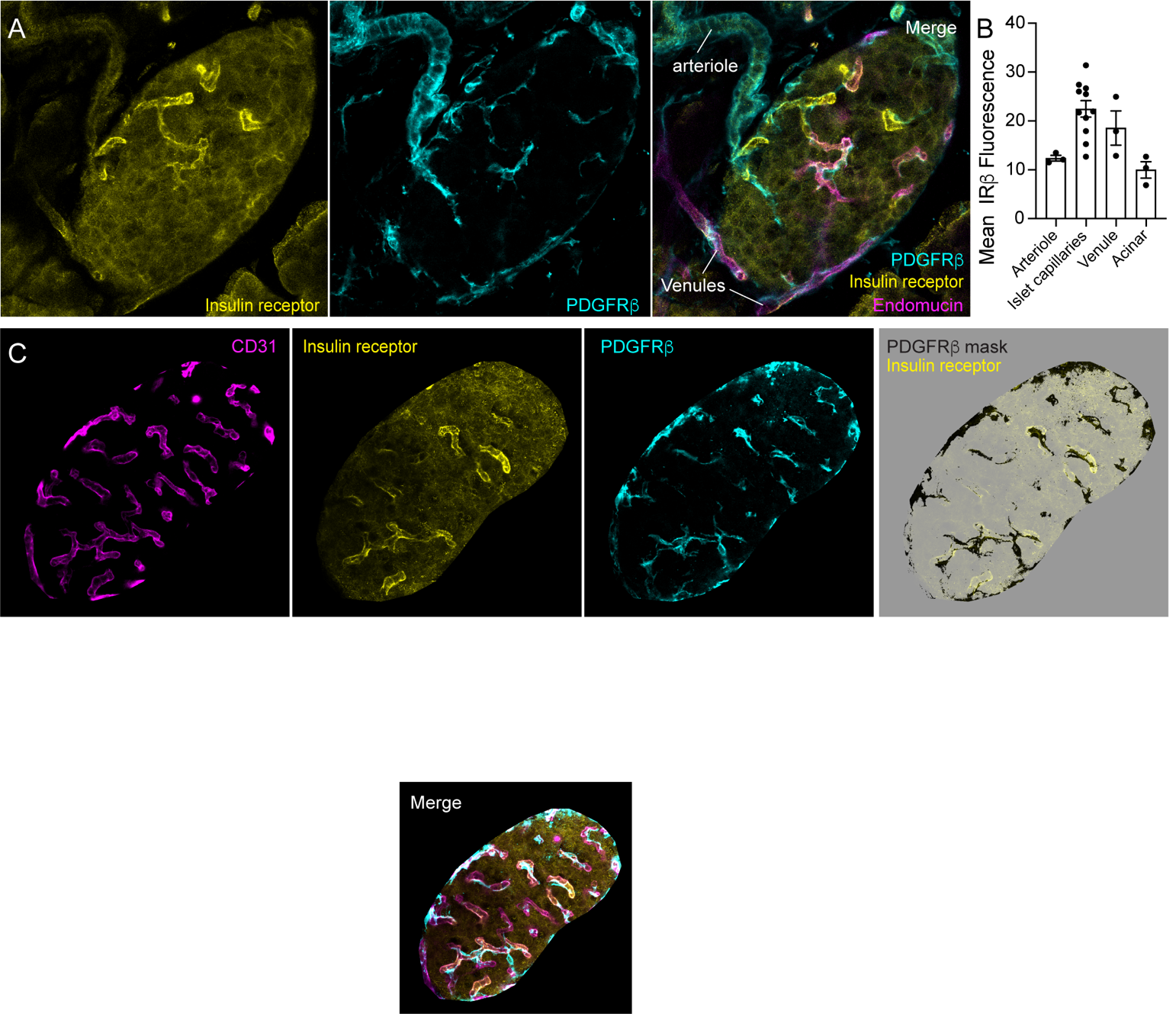

**Supplementary Figure 2. Association between PDGFRβ-expressing pericytes and islet capillaries expressing insulin receptors.** (A,C) Only a subset of the islet microvasculature expresses insulin receptors (antibody against its β subunit was used; yellow). Those are vessels covered with PDGFRβ-expressing pericytes (cyan) inside pancreatic islets. Endomucin was used to visualize venous endothelial cells (magenta). (B) Mean fluorescence intensity of insulin receptor β signal is higher in islet capillaries and venules (endomucin positive) than in the islet arteriole or surrounding acinar tissue. Different regions of interest were placed around these structures in the islet shown in (A). (C) Based on PDGFRβ signal (cyan), we created a mask (black) that we applied on top of the insulin receptor channel (yellow). Intra-islet capillaries (visualized with CD31; magenta) that express insulin receptor are those with highest PDGFRβ coverage on their surface (right panel: merge image of PDGFRβ mask and insulin receptor channel (with 40% opacity)). Only the islet is shown (regions outside were cleared out).

**Supplementary Figure 3**

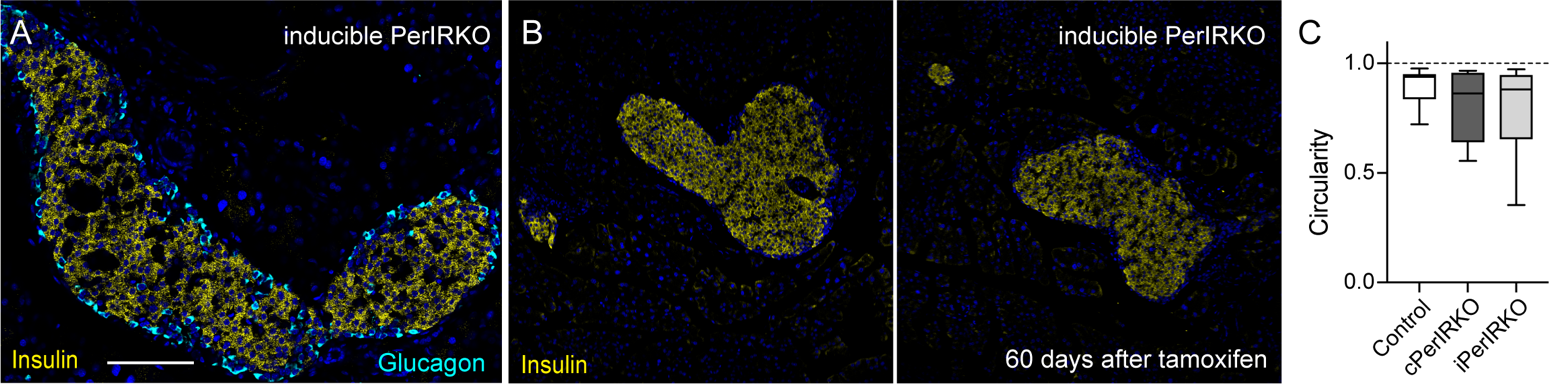

**Supplementary Figure 3. Some islets lose their circular shapes as they increase in area in iPerIRKO mice.** (A,B) Different islets are shown in pancreatic sections from inducible PerIRKO mice immunostained for insulin (yellow) and glucagon (cyan; A) 60 days after tamoxifen treatment. Different islets seem to have merged with each other. Scale bar = 100 μm (A). (C) Quantification of the circularity of islets in control and PerIRKO mouse models using ImageJ shape descriptors.

**Supplementary Figure 4**
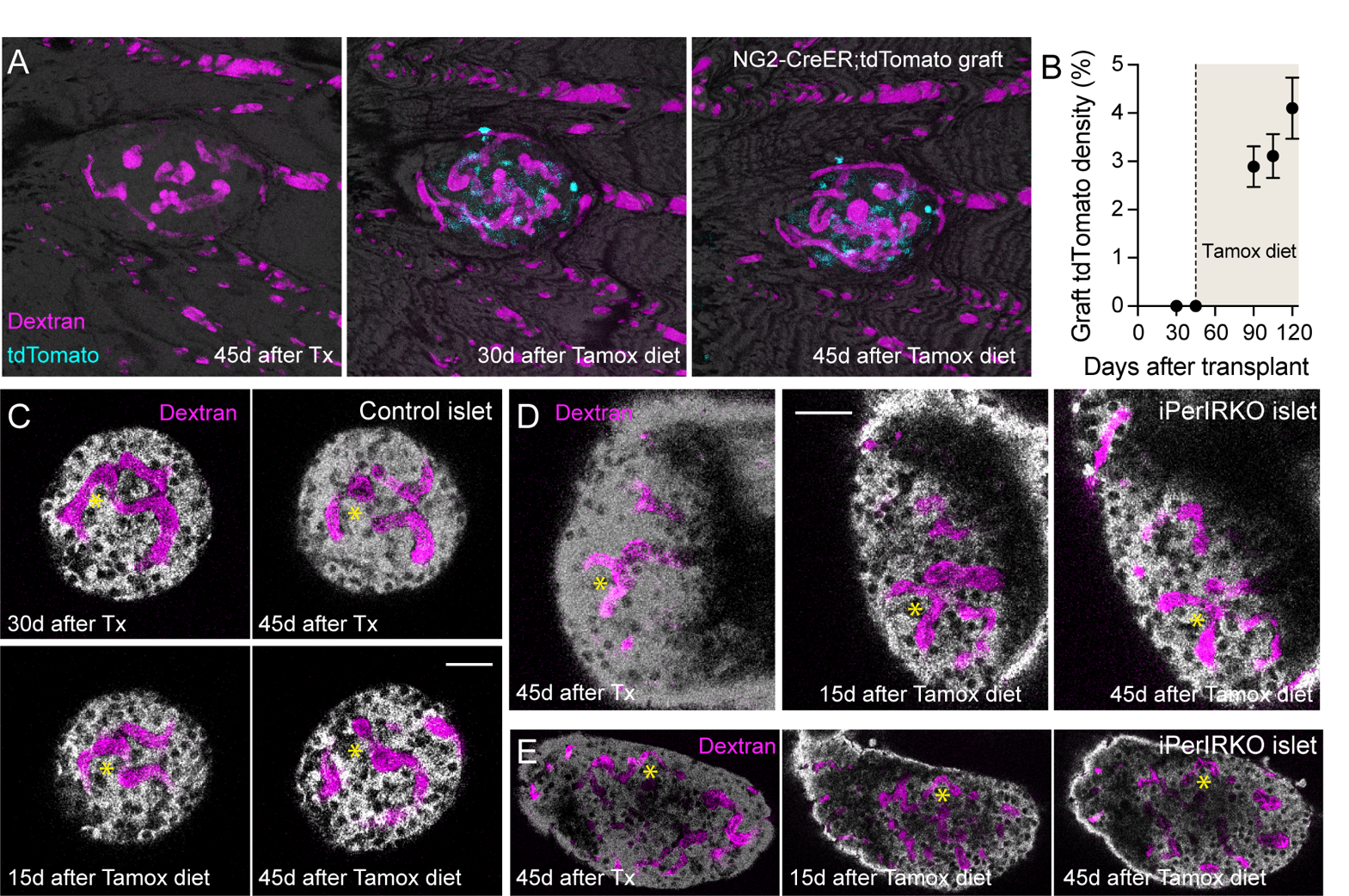

**Supplementary Figure 4. Longitudinal imaging of intraocular islet grafts from inducible PerIRKO and control mice.** (A) Z-projection of confocal images of an intraocular islet from a NG2-CreER;tdTomato^fl/fl^ mouse 45 days after transplantation (left panel) and then 30 (middle panel) or 45 days (right panel) after placing mice on a tamoxifen-containing diet to induce tdTomato (cyan) expression in pericytes. Blood vessels were labeled with FITC-dextran (150 kDa; magenta). (B) Quantification of tdTomato-positive area divided by total graft area at different time points before and after tamoxifen diet (shaded area). (C-E) Confocal images of a control (C) and two iPerIRKO (D,E) islet grafts at different time points after transplantation (Tx) and after starting tamoxifen (tamox) diet feeding. Vessels are shown in magenta (dextran) and endocrine cells visualized with the backscatter signal (gray). Yellow * is used to indicate the same structure in each islet graft during different imaging sessions to ensure the same confocal plane is being visualized. Scale bars = 20 μm (C,D).

**Supplementary Movies**

**Supplementary Movie 1. Recording vasomotor responses to norepinephrine and high glucose in a living pancreatic slice from a control mouse.** Series of Z-projections of confocal images of lectin-labeled blood vessels in a control islet (IR^fl/fl^ male mouse treated with tamoxifen) before and during stimulation with norepinephrine (NE; 20 μM) followed by washout period with 3 mM glucose (3G) and then stimulation with 16 mM glucose (16G). Grayscale images were inverted for better contrast. Movie speed is of 40 fps. Notice constriction of a subset of islet capillaries and arteriole in response to NE and dilation triggered by 16G.

**Supplementary Movie 2. Recording islet vasomotor responses to high glucose in a living pancreatic slice from a control mouse.** Series of Z-projections of confocal images of lectin-labeled blood vessels in a control islet (IR^fl/fl^ male mouse treated with tamoxifen) before and during stimulation with 16 mM glucose (16G). Grayscale images were inverted for better contrast. Movie speed is of 40 fps. Notice dilation of islet capillaries in response to 16G stimulation.

**Supplementary Movie 3. Recording islet vasomotor responses to norepinephrine in a living pancreatic slice from an iPerIRKO mouse.** Series of Z-projections of confocal images of lectin-labeled blood vessels in an islet from an iPerIRKO female mouse (60 days after tamoxifen) before and during stimulation with norepinephrine (NE; 20 μM). Grayscale images were inverted for better contrast. Movie speed is of 40 fps. Notice constriction of a subset of islet capillaries in response to NE stimulation.

**Supplementary Movie 4. Recording islet vasomotor responses to norepinephrine in a living pancreatic slice from iPerIRKO mouse.** Series of Z-projections of confocal images of lectin-labeled blood vessels in an islet from an iPerIRKO female mouse (60 days after tamoxifen) before and during stimulation with norepinephrine (NE; 20 μM). Grayscale images were inverted for better contrast. Movie speed is of 40 fps. Notice vessels in the middle of the islet with a “spring” shape which move towards each other in response to NE stimulation.

**Supplementary Movie 5. Recording islet vasomotor responses to high glucose in a living pancreatic slice from iPerIRKO mouse.** Series of Z-projections of confocal images of lectin-labeled blood vessels in an islet from an iPerIRKO female mouse (60 days after tamoxifen) before and during stimulation with 16 mM glucose solution. Grayscale images were inverted for better contrast. Movie speed is of 40 fps. Notice vessels with “spring” shape in the middle of the islet which do not respond to high glucose, while capillaries at the islet border dilate in the presence of 16G.
